# A logical model of homology for comparative biology

**DOI:** 10.1101/588822

**Authors:** Paula M. Mabee, James P. Balhoff, Wasila M. Dahdul, Hilmar Lapp, Christopher J. Mungall, Todd J. Vision

## Abstract

There is a growing body of research on the evolution of anatomy in a wide variety of organisms. Discoveries in this field could be greatly accelerated by computational methods and resources that enable these findings to be compared across different studies and different organisms and linked with the genes responsible for anatomical modifications. Homology is a key concept in comparative anatomy; two important types are historical homology (the similarity of organisms due to common ancestry) and serial homology (the similarity of repeated structures within an organism). We explored how to most effectively represent historical and serial homology across anatomical structures to facilitate computational reasoning. We assembled a collection of homology assertions from the literature with a set of taxon phenotypes for vertebrate fins and limbs from the Phenoscape Knowledgebase (KB). Using six competency questions, we evaluated the reasoning ramifications of two logical models: the Reciprocal Existential Axioms Homology Model (REA) and the Ancestral Value Axioms Homology Model (AVA). Both models returned the user-expected results for all but one historical homology query and all serial homology queries. Additionally, for each competency question, the AVA model returns the search term and any subtypes. We identify some challenges of implementing complete homology queries due to limitations of OWL reasoning. This work lays the foundation for homology reasoning to be incorporated into other ontology-based tools, such as those that enable synthetic supermatrix construction and candidate gene discovery.

## Introduction

Distinguishing homology, i.e., similarity due to inheritance from a common ancestor, from similarities that arise independently, is the foundation of the comparative approach that is applied across many different fields of biology. Comparative genomics, for instance, has led to the identification of homologous patterns of gene activity and regulation that have been conserved over hundreds of millions of years of evolution. This has been aided considerably by computer-based analysis, which is enabled by the standardization of genomic data. The longer tradition of comparative anatomy has also revealed extensive conservation, with the homologies between the jaw bones of fishes and the inner ear bones of mammals as a quintessential example. The complexity of anatomical data, however, has been an impediment to standardization and computation, and many of the critical tasks rely on manual inspection of the data and human judgment [1]. Advances in this area have been made using semantic reasoning, but these have not explicitly incorporated nor evaluated homology reasoning. Here we formalize the biological expectations for homology reasoning and evaluate the consequences of applying formal homology relationships between anatomical structures in an anatomy ontology.

Anatomy ontologies are formal graph representations of anatomical structures and the relationships among them. They provide the foundation for computational analyses of comparative anatomy data that are semantically aware. By aggregating expert knowledge of different anatomical structures and organisms, they are a key resource for comparative analysis. Anatomy ontologies exist today that can connect the anatomical features and linked data from millions of biological species.

The Phenoscape project (www.phenoscape.org) has been working to demonstrate the value of a semantic approach through the development of multi-species anatomy ontologies [2–5] and other required resources including taxonomy ontologies [6], annotation tools [7,8], and a knowledgebase to hold these structured data [9]. Phenoscape has annotated >22,000 anatomical character states to >5,000 vertebrates from the comparative evolutionary literature and integrated the resulting more than half a million taxon phenotypes with approximately 400K gene phenotype annotations from model organisms (zebrafish, mouse, *Xenopus* and human) into its knowledgebase. Using ontology-based reasoning to integrate taxon and gene phenotypes, the team has demonstrated the discovery of candidate genes underlying evolutionarily novel phenotypes [10] and the use of semantic similarity to discover evolutionary variation related to gene phenotypes [11].

The anatomy ontologies and reasoning capabilities of the Phenoscape Knowledgebase (hereafter, the KB) provide the core framework for automatically extracting the basic data desired at the outset of a comparative anatomical study, namely all of the published data for a set of anatomical structures in a focal taxon. Although a researcher might use these data in a number of different ways, the data required will generally be a matrix of taxa and anatomical phenotypes. An illustration of such reasoning is provided by the Ontotrace tool, which can directly extract or infer the presence or absence of anatomical entities across all studies in the KB for a user-specified set of anatomical entities and taxa; these can then be aggregated into an aligned matrix for downstream analysis [12,13].

To date, reasoning performed in the KB using anatomy ontologies has not explicitly incorporated homology. The present work attempts to address that deficiency, but it is not a straightforward task, in part because homology can be defined in numerous ways. The literature is replete with continuing discussions about types of homology, levels of homology, and how to distinguish homology from homoplasy, i.e., similarity that is not due to common ancestry (convergent or parallel evolution) [14–25]. The similarity of features that are descended from a common ancestor is typically referred to as ‘historical homology’ [18]; the homologies between the jaw bones of fishes and the inner ear bones of mammals are a quintessential example [26]. ‘Serial homology’, a type of iterative homology, is the historical and developmental relationship among segmented or, more generally, iterated, structures within an organism, *e.g*., across the various appendages of crustacea, the vertebrae of vertebrates, and the arms of a starfish [16]. Despite the volume of literature on homology, an explicit mapping from the biological understanding of these types of homology relationships to their downstream logical consequences has not been made. For instance, given an assertion of serial or historical homology between two anatomical structures, how should that knowledge be logically propagated to their parts, subtypes, or developmental precursors? Having explicit logic that mirrors biologists’ expectations but that can also be employed computationally would enable computationally-assisted discoveries in comparative biology only limited by the scale of semantically-described biodiversity data this is available.

### How to accommodate homology within ontologies?

A number of approaches to using information about homology in relating anatomical entities or phenotypes in ontologies have been proposed or implemented. Early ideas for incorporating homology grew out of the effort to expand the taxonomic scope of anatomy ontologies beyond the single species, typically model organisms, for which many of them had been designed [27]. It was argued that because similarity of phenotype frequently owes to the continuity of inherited information, *i.e.*, homology, that it must be accommodated in any attempt to create multi-species anatomy ontologies [27]. Several approaches were considered, one of which was to represent homologs with different names as synonyms of a single anatomical entity [28]. For example, the series of bones located along the midline between the skull and dorsal fin in different fish species is referred to by different names (‘supraneural’ and ‘predorsal’). Given the homology of these series across all fishes [29], they are represented in the Uberon anatomy ontology by a single concept under the term ‘supraneural’ with the exact synonym ‘predorsal’. Although this representation might suffice for structures with strong evidence of homology, it does not accommodate differently named structures with very different structural, developmental, and positional relationships. For example, the stapes, an inner ear bone in mammals, is the undisputed homolog of the hyomandibula, a jaw bone in fishes. If these terms were synonymized, many specific relationships would need to be generalized for the ontology to accurately represent both the stapes and hyomandibula. Synonymizing also does not suffice in cases where the homology is uncertain. For example, the ‘alular digit’ (*i.e.*, the first digit in bird wings) is considered by many, on the basis of paleontological evidence [30], to be a homolog of the first digit in other vertebrates (*i.e.*, ‘manual digit 1’). However, these are typically not considered homologous on the basis of developmental data [31,32].

Another proposed approach was to represent hypotheses for the homology of anatomical entities outside of formalized ontologies; the ontology itself could remain homology-neutral [28], also see [1–3]. Anatomical entities would be defined on the basis of spatio-structural properties that would allow their unambiguous identification and re-identification exclusively on the basis of anatomy [1]. This approach is further justified by the fact that at least some homology hypotheses are too weak or controversial to be embedded in the ontology in the same way as the hardened knowledge concerning the types and parts of anatomical structures. This approach, *i.e.*, to capture hypotheses of homology independently of structural and functional information, was supported by Haendel *et al.* [33], who further proposed a new relationship, *homologous_to*, to be included in the OBO Relations Ontology. This relationship was further defined and formalized, along with *not_homologous_to*, by Travillian *et al.* [34], who implemented them in the Vertebrate Bridging Ontology (VBO). The VBO was introduced to enable the transfer of information about homologous anatomical structures between species-specific anatomical ontologies, and a beta version was integrated into the Experimental Factor Ontology [35] to support cross-species comparisons of orthologous genes in homologous tissues through the Gene Expression Atlas interface.

The Bgee initiative (bgee.org) led computational work to use homology relations to align anatomical entities between species-specific anatomy ontologies to enable comparisons of gene expression patterns between species [25,36,37]. These authors designed an algorithm, implemented in the software Homolonto [36], to create new relationships between anatomical ontologies, and an ontology to clarify homology-related concepts, a homology ontology (HOM) [38]. They later developed the vertebrate Homologous Organs Groups ontology (vHOG), a multispecies anatomical ontology for vertebrates based on the homologous organ systems used in the Bgee database of gene expression evolution [39]. vHOG describes structures with historical homology relations between model vertebrate species, and includes manually reviewed mappings to species-specific anatomical ontologies (no homology hypothesis is stated within the ontology itself) [39].

Currently, in multispecies anatomy ontologies (*e.g.*, Uberon [5]; TAO [2]; VSAO [3]), the definitions of classes focus on some re-identifiable property or properties that members of the classes have in common; they do not include criteria of homology. Most often these properties involve structural criteria, but developmental and functional ones are employed, too. A class such as ‘endochondral bone’ (UBERON:0002513) reflects the developmental similarity of its subtypes, ‘long bone’ (UBERON:0002495) reflects structural similarity, and ‘eye’ (UBERON:0000970) reflects functional similarity. This pluralistic approach reflects the multiple ways that comparative morphologists understand and group anatomical structures. By not imposing homology on the ontology, one might argue that broader possibilities for data discovery are enabled. Simply, searches for similar anatomical structures are not constrained by the special similarity owing to shared evolutionary descent.

That notwithstanding, for many classes in multi-species anatomy ontologies, and therefore Uberon [5], homology is implicit in their semantics, as evidenced by how they are applied in practice. For example, the most proximal bone of the forelimb/arm in vertebrates, including humans, is named the ‘humerus’. The humerus is considered a historically homologous bone across vertebrates, and the single label for this bone in Uberon, i.e., ‘humerus’, signifies homology in this case. Expert curators use this term across vertebrates without restricting the semantics to different taxonomic groups. Homology, in fact, is thus woven into the names of many if not most anatomical structures [37], and a multi-species anatomy ontology therefore cannot be characterized as ‘homology-free’ [11].

Finally, some arguments have been made to bake homology into the ontology [40], embodying the knowledge derived about character state homology from phylogenetic trees. Although this is a solution for well supported hypotheses of homology, for others that are disputed, it is not. Further, because proposals of homology are tested by concordance with phylogeny, to the extent that phylogenetic hypotheses themselves are in flux, hypotheses of homology are as well.

Our goal with this study is to understand the ramifications of different ways of representing historical and serial homology for anatomical entities as a set of ontology axioms. We first describe two alternative models which differ in their requirements for logical expressivity. We then assess the implications of these differing models when applied to the classic example of homologies between fish fins and tetrapod limbs [41]. The assessment is guided by six competency questions that aim to capture reasonable user expectations for how an assertion of homology between two anatomical entities should affect subsequent reasoning.

We close with a discussion of implementation of one homology model within the Phenoscape Knowledgebase, which must balance logical expressivity with the practicalities of scalability across a large dataset.

## Methods

### Logical models of homology assertions

To be used by an OWL reasoner, and thus have an effect on reasoner-driven query resolution, each homology assertion must be translated into OWL axioms. For an explanation of the types of axioms which can be stated within OWL ontologies, see [42]. Modeling in OWL frequently involves a tradeoff between expressivity and scalability. In fact, the OWL language provides three “profiles”—subsets of the language which omit the ability to make certain kinds of statements—which are known to be amenable to more scalable reasoning algorithms [43]. Because each profile omits different capabilities, each is better suited to particular kinds of modeling tasks. The OWL EL profile is frequently used with complex biomedical terminologies, such as large anatomy ontologies. The availability of efficient EL reasoners such as ELK has been been crucial to the application of OWL in the development of ontologies like the Gene Ontology and Uberon [44].

Here, we introduce two alternative logical models for homology: Reciprocal Existential Axioms (REA) and Ancestral Value Axioms (AVA) (**Table 1**). REA is designed to fit within the OWL EL profile. AVA, on the other hand, provides semantics which may potentially be a more exact fit to user expectations of homology but requires reasoning capabilities which are not part of the scalable OWL EL profile.

**Table 1.**
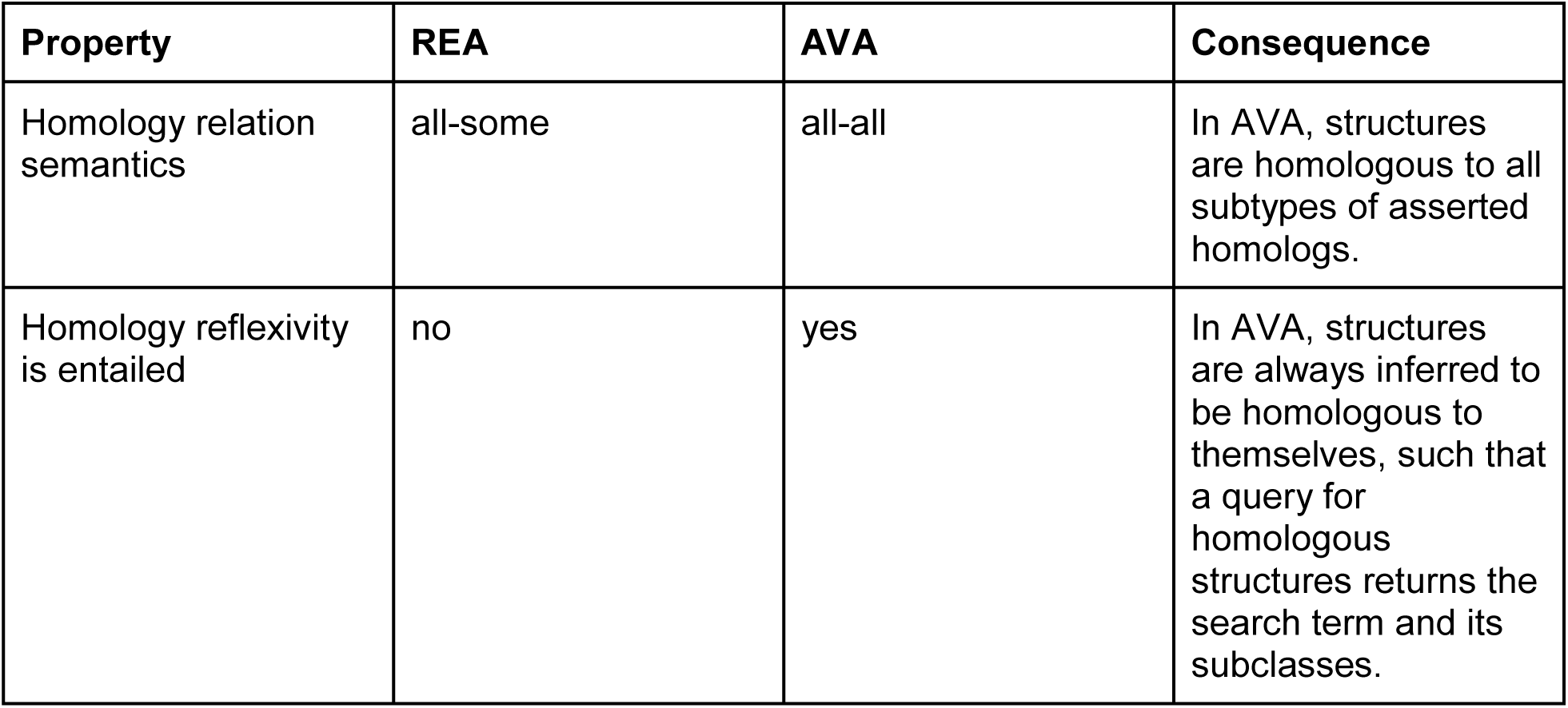
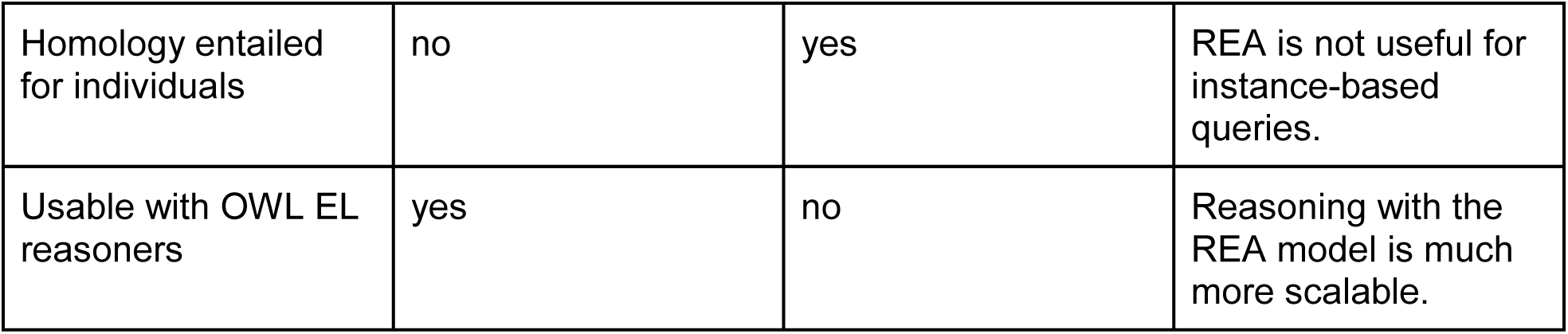
Comparison of the REA and AVA models.

| Property | REA | AVA | Consequence |
| --- | --- | --- | --- |
| Homology relation semantics | all-some | all-all | In AVA, structures are homologous to all subtypes of asserted homologs. |
| Homology reflexivity is entailed | no | yes | In AVA, structures are always inferred to be homologous to themselves, such that a query for homologous structures returns the search term and its subclasses. |
| Homology entailed for individuals | no | yes | REA is not useful for instance-based queries. |
| Usable with OWL EL reasoners | yes | no | Reasoning with the REA model is much more scalable. |

As its name states, REA models a homology annotation as a reciprocal pair of axioms. Each of the two structures is given a subclass relationship to an existential property restriction using the other structure (*i.e.*, every instance of the first structure is homologous to some instance of the second structure):

~~~
‘pectoral fin’ SubClassOf (homologous_to some ‘forelimb’)
‘forelimb’ SubClassOf (homologous_to some ‘pectoral fin’)
~~~

This model has the advantage that it can be rapidly classified and queried using efficient OWL reasoners implementing the OWL EL profile. REA only enables querying via class expressions, though in the Phenoscape KB this is the most common query approach and thus not a limitation. For example, with the REA model, one can query for all subclasses of the class expression `homologous_to some ‘forelimb’`, and all instances of this expression. However, as a consequence of the REA model, given a particular instance of ‘forelimb’, one cannot find other individuals inferred to be homologous to it, because the respective axiom asserts only that (*all*) instances of ‘forelimb’ are homologous to *some* instance of ‘pectoral fin’, not to all instances of it. This is sometimes referred to as “*all-some*” semantics.

The AVA model introduces, for each homology annotation, an OWL individual (*i.e*., an instance) that represents the ancestral structure from which all instances of the two classes of homologous structures are descended:

~~~
‘pectoral fin’ SubClassOf historical_homology_member_of value <pectoral_fin_forelimb_ancestor>
‘forelimb’ SubClassOf historical_homology_member_of value <pectoral_fin_forelimb_ancestor>
~~~

Two additional property axioms provide entailment of the needed `homologous_to` relationship:

~~~
historical_homology_member_of InverseOf: has_historical_homology_member
historical_homology_member_of ⃭ has_historical_homology_member ⃭ homologous_to
~~~

The result of these axioms is an “*all-all*” semantics, as opposed to the all-some semantics of REA. That is, this model entails for any two instances of ‘pectoral fin’ and ‘forelimb’ that they stand in a homologous_to relationship to each other. Properties with all-all semantics are exceedingly rare, at least in most ontologies encoding biological knowledge domains, because most biologically important relationships can be universally asserted only in one direction. For example, the part_of relationship common in anatomy ontologies (such as Uberon) holds between two anatomical entities as `A SubClassOf (part_of some B)`, as in `humerus SubClassOf (part_of some forelimb)`. The all-some semantics entails that a given instance of humerus is part of one specific forelimb; it is not part of every instance of forelimb. The stronger *all-all* semantics provided by AVA may more closely match the expectations of a user who asserts historical homology between two anatomical entities. However, the logical expressivity needed for reasoning with AVA requires features, such as inverse property axioms, that are outside of the OWL EL profile, which in practice makes this model much less scalable.

With either model, one can assert homology between structures in a way that is taxonomically more restrictive than implied by the way that the corresponding anatomy ontology terms are defined. To account for such restrictions, we substitute the anatomical entities A and B with taxon-based subset expressions. More formally, if specifically entity A occurring in taxon X is homologous to entity B occurring in taxon Y, we substitute A with the class expression “A and in_taxon some X” (*i.e.*, those instances of A that are in some instance of taxon X), and B with “B and in_taxon some Y” (here using the REA model):

~~~
(A and in_taxon some X) SubClassOf (homologous_to some (B and in_taxon some Y))
(B and in_taxon some Y) SubClassOf (homologous_to some (A and in_taxon some X))
~~~

Here X and Y are terms from a taxonomy ontology, *e.g.* Vertebrate Taxonomy Ontology (VTO) [6]. The `in_taxon` relation (RO:0002162) is used throughout Uberon to specify “taxonomic constraints” on anatomical concepts. It is used primarily for automated quality control of annotations and for consistency checking when merging independently developed anatomy ontologies into Uberon.

To relate two anatomical terms as homologous, we used the relation ‘in historical homology relationship with’ (RO:HOM0000007) (above referenced as ‘homologous_to’) from the OBO Relations Ontology (RO; http://purl.obolibrary.org/obo/ro.owl), which is defined as: “Homology that is defined by common descent.” Serially homologous structures were related using the relation ‘in serial homology relationship with’ (RO:HOM0000027), which is defined as: “Iterative homology that involves structures arranged along the main body axis.” These relations are derived from the Homology Ontology (HOM; http://purl.obolibrary.org/obo/hom.owl), which contains 66 classes representing concepts related to organismal similarity, including homology and homoplasy. Classes of homology from this ontology are mirrored as object properties within RO, providing the relationships needed to assert historical or serial homology between anatomical structures. For testing the AVA model, we used locally defined properties for ‘historical_homology_member_of’ and ‘has_historical_homology_member’ (and corresponding relations for serial homology), as these are not currently defined in the Relation Ontology.

### Biological expectations for homology reasoning

To evaluate the consequences of applying a formal homology relationship between anatomical structures, we establish specific user expectations in the form of competency questions [45,46] for the results of a description logic (DL) query of our demonstration ontology. Competency questions are a set of questions that an ontology must answer using the knowledge represented by its axioms [45,46]. A DL query is an OWL expression logically describing a class for which we want to find its subclasses or instances (for example). Our competency question expressions are modeled using the relations composing the EQ phenotypes (anatomical entity (E) and a quality (Q) [47,48]) in the test dataset, but the qualities themselves do not play a role in the homology reasoning. Put another way, `inheres_in some (homologous_to some ‘pectoral fin’)` would subsume any phenotype instances referring to homologs of the pectoral fin. We focus on expectations about how homology is inferred across the broader ontology graph in which the anatomical structures are embedded, such as “to what degree is an assertion of homology propagated to other structural, positional, and developmental, relationships?” and “to what degree are homology relationships transitive?”

To our knowledge, this is the first attempt to formalize expectations for homology reasoning in a general manner suitable for evaluating a semantic model. These expectations are framed from the standpoint of a hypothetical user, a comparative evolutionary anatomist who is well-versed in the data that pertain to homology of the structures under consideration. The expectations of this persona guide the general way in which the logic of a homology relationship between two structures propagates beyond them to their parts, types, developmental precursors, developmental products, and other homologs. While some expectations are clear, and would be so to any biologist (*e.g.*, that the parts of homologous structures are not necessarily homologs), others might be debatable. In these cases we take a conservative approach wherever homology reasoning might lead to incorrect inferences. For example, some might desire homology reasoning to lead to the conclusion that the developmental precursors of homologous structures are themselves homologs. Given the evidence that this is incorrect in some cases (i.e., developmental precursors of homologs are not themselves homologous), extending the homology reasoning to developmental precursors is not permitted. Thus, homology reasoning in our models involves only subsumption (‘is_a’) relations, and reasoning to other relationship types (i.e., develops_from, part_of) would need to be executed via property chain reasoning not employed in our homology models. Overall, we take the approach to formulate general expectations for inferred results only for relationships for which the propagation of homology should generally be biologically correct or accepted.

The expectations that are generally applicable to any structure for historical and serial homology queries are the following: In the case of historical homology, queries for historical homologs of a structure are expected to return historical homologs, subtypes of historical homologs, historical homologs of the superclass and its subtype(s), and taxonomically restricted results. Further, queries for historical homologs of a structure are not expected to return parts of the historical homolog, serial homologs, developmental products (i.e., structures that later develop from the historical homolog), or developmental precursors of the historical homolog.

In the case of serial homology, queries for serial homologs of a structure are expected to return serial homologs and subtypes of serial homologs, and serial homologs of the superclass and its subtype(s). Further, queries for serial homologs of a structure are not expected to return parts of the serial homolog, historical homologs of the serial homologs, historical homologs of the superclass and subtype(s), developmental precursors of the serial homolog, or developmental products or developmental precursors of the serial homolog.

### Competency questions

We created seven competency questions to test the expectations of our biologist persona for the results of a query using historical or serial homology.

#### Competency question 1

Our biologist persona expects a phenotype query for historical homologs of **‘pectoral fin’** to return phenotypes for its homolog ‘forelimb’ and its homolog’s subtype ‘forelimb wing’, as illustrated in **Figure 1**. They do not expect the query to return phenotypes for parts of the homolog, such as ‘humerus’, nor phenotypes for serial homologs of the homolog (‘hindlimb’). Further, they do not expect it to return phenotypes for the homolog’s developmental precursor ‘forelimb bud’.

**Figure 1.**
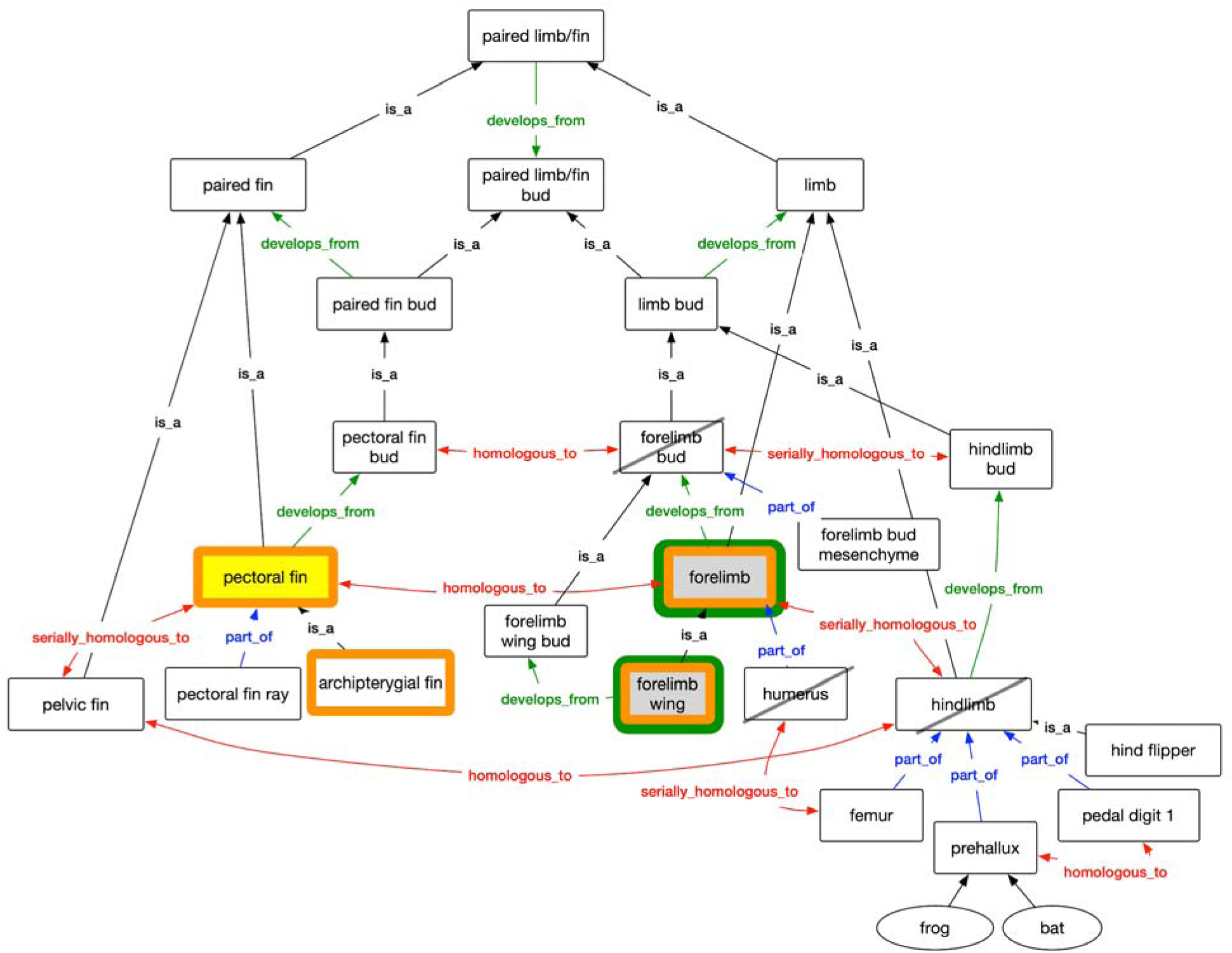
Expectations for competency question 1, a query for historical homology of class ‘pectoral fin’ (bright yellow fill). The persona expected the classes ‘forelimb’ and ‘forelimb wing’ (grey fill), to be returned. Unexpected classes (black slash) would include parts (‘humerus’), serial homologs (‘hindlimb’), and developmental precursors (‘forelimb bud’). Both REA (green outline) and AVA (orange outline) queries returned all the results that were expected; In addition, AVA returned the search term (‘pectoral fin’) and its subtype (‘archipterygial fin; orange border’).

#### Competency question 2

Our persona expects a phenotype query for historical homologs of ‘forelimb wing’ to return phenotypes for ‘forelimb’, ‘pectoral fin’, and subclasses of ‘pectoral fin’ such as ‘archipterygial fin’ (**Figure 2**). They do not expect to be returned structures including parts (e.g., ‘humerus’ or ‘pectoral fin ray’) of the homologs, serial homologs (‘hindlimb’ or ‘pelvic fin’) of the homologs, and developmental precursors (‘forelimb bud’ or ‘pectoral fin bud’) of the homologs.

**Figure 2.**
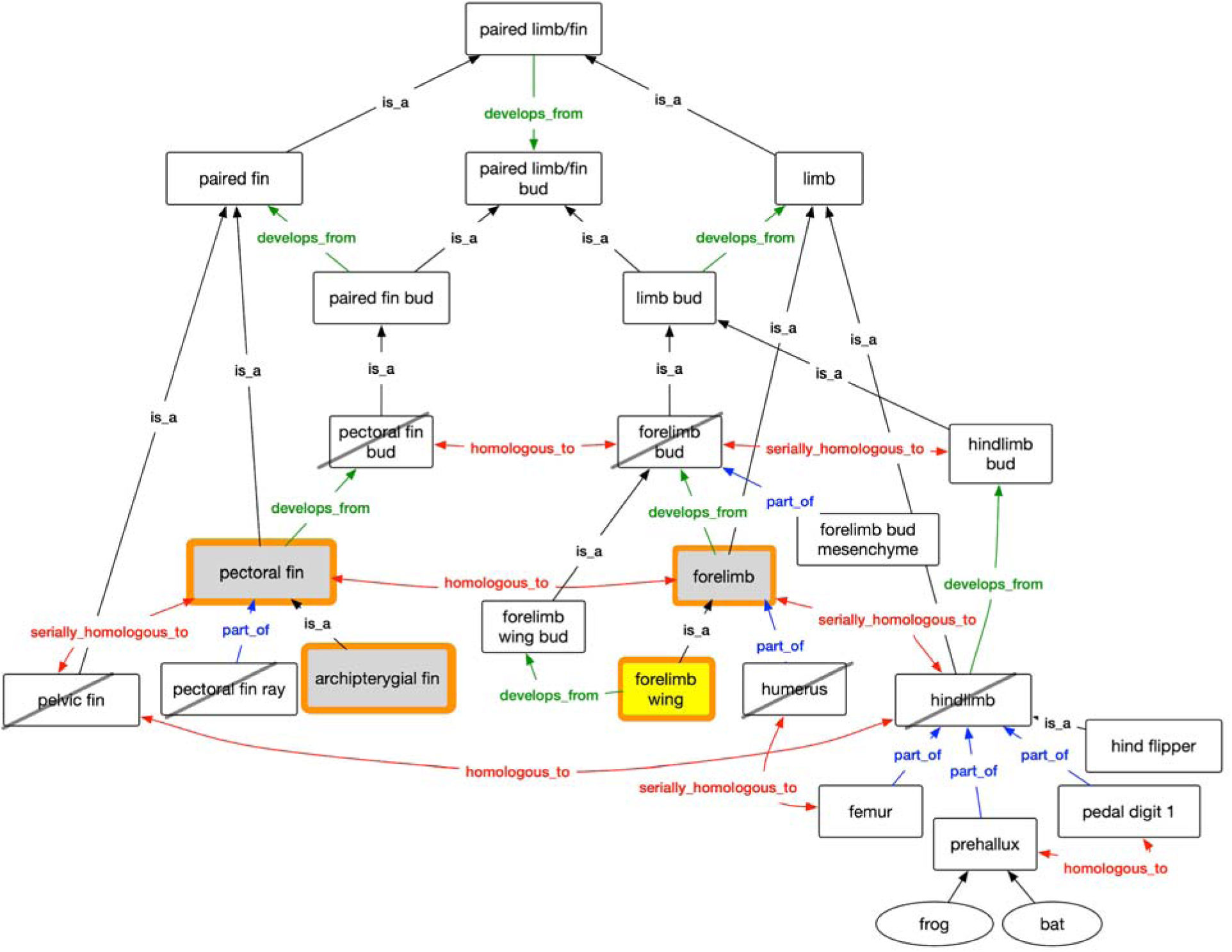
Expectations for competency question 2, a query for historical homology of class ‘forelimb wing’ (bright yellow fill). Our persona expected ‘forelimb’, ‘pectoral fin’ and ‘archipterygial fin’ (grey fill) to be returned. Structures not expected (black slash) include parts (‘humerus’, ‘pectoral fin ray’), serial homologs (‘hindlimb’, ‘pelvic fin’), and developmental precursors (‘forelimb bud’, ‘pectoral fin bud’). The results of the AVA query (orange outline) included expected structures in addition to the search term. In REA, no results were returned.

#### Competency question 3

Our persona expects a query for historical homologs of **‘pectoral fin bud’** to return phenotypes for ‘forelimb bud’ and its subtype ‘forelimb wing bud’ (**Figure 3**). They do not expect it to return phenotypes to parts (‘forelimb bud mesenchyme’), structures that form later in the course of development (‘forelimb’), or serial homologs (‘hindlimb bud’) of the historical homologs.

**Figure 3.**
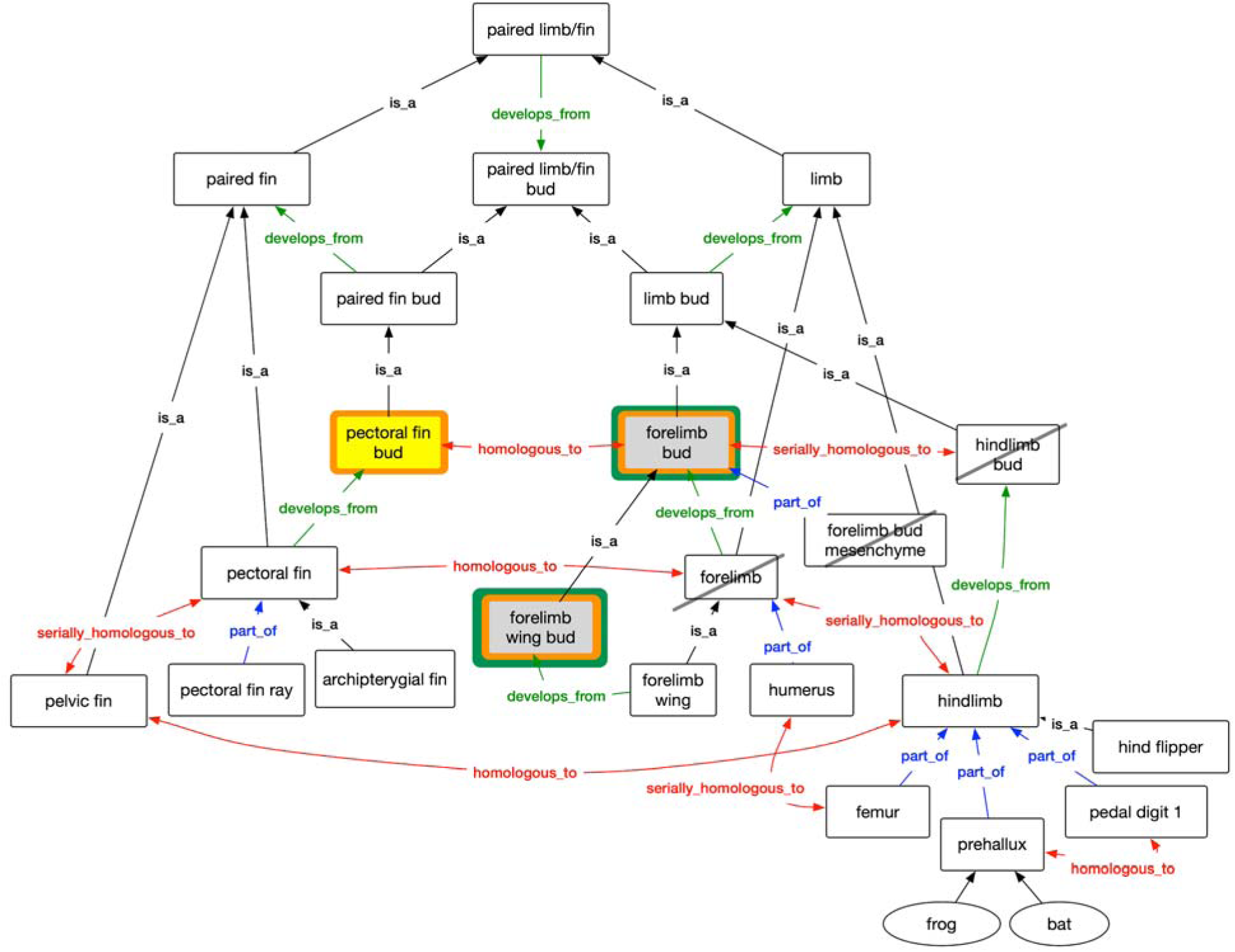
Expectations for competency question 3, a query for historical homology of class ‘pectoral fin bud’ (bright yellow fill). Our persona expected ‘forelimb bud’ and ‘forelimb wing bud’ (grey fill) to be returned. Structures not expected (black slash) include parts (‘forelimb bud mesenchyme’), serial homologs (‘hindlimb bud’), and structures that later develop (‘forelimb’). Both REA (green outline) and AVA (orange outline) queries returned all the expected results; In addition, AVA returned the search term (‘pectoral fin bud’).

#### Competency question 4

In cases where the homology statement selectively applies to a subset of taxa that possess the anatomical structure, though other taxa may ostensibly possess it as well, our persona expects results for only a restricted set of taxa. They expect a query for the historical homologs of ‘**pedal digit 1**’ to return ‘prehallux’ phenotypes for only anurans (frogs) and not ‘prehallux’ phenotypes for mammals (bats in our example)^1^ (**Figure 4**). This is because the homology relationship is specific to Anura and pedal digit 1.

**Figure 4.**
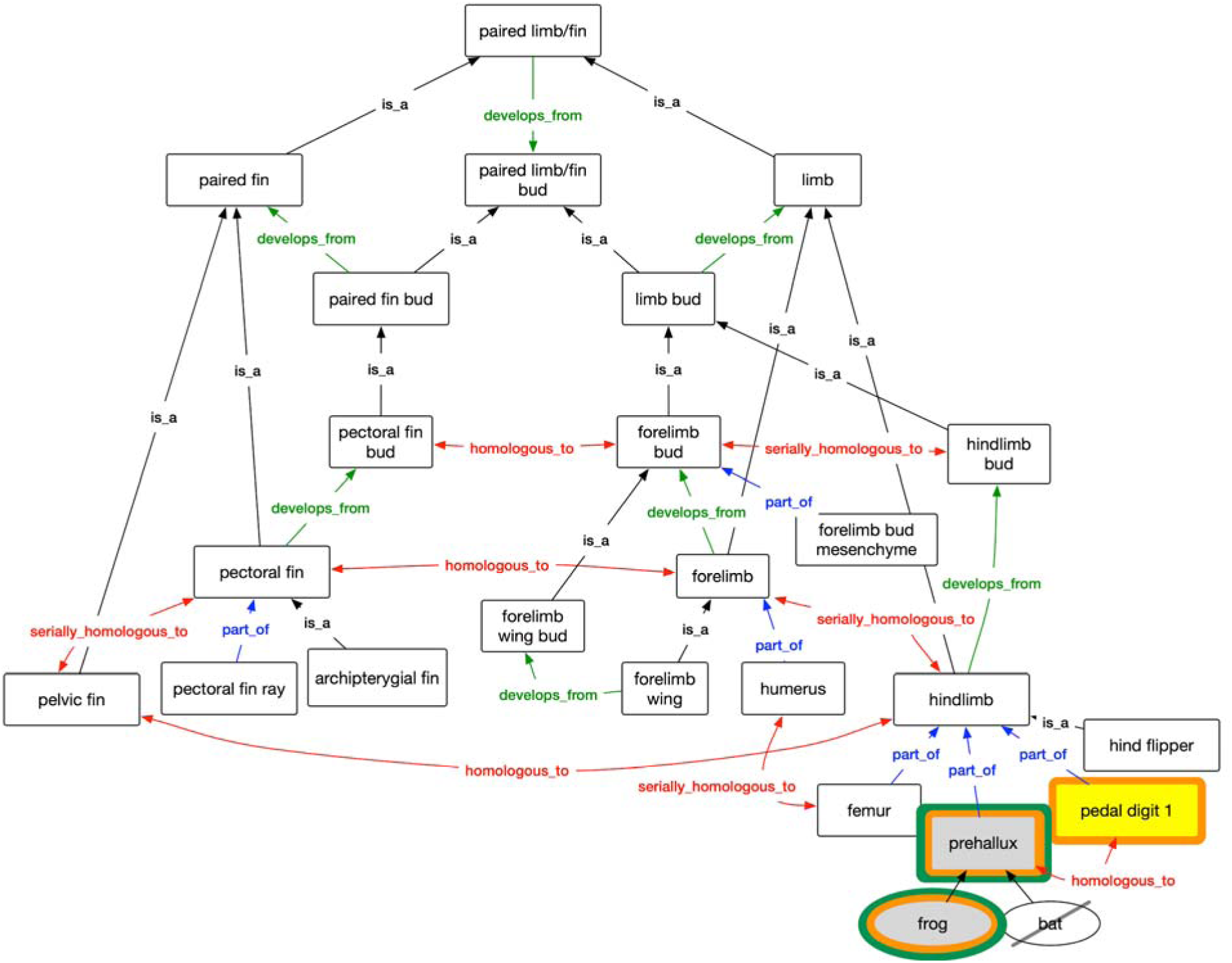
Expectations for competency question 4, a query for historical homology of class ‘pedal digit 1’ (bright yellow fill). Our persona expected phenotypes for ‘prehallux’ to be returned for only the frog (grey fill). Phenotypes for ‘prehallux’ that are associated with mammals (bat, **Table 3**), are not expected (black slash). Both REA (green outline) and AVA (orange outline) returned the expected results; AVA additionally returned ‘pedal digit 1’.

#### Competency question 5

From a query on serial homologs of **‘hindlimb’**, our persona expects to find phenotypes for ‘forelimb’ and its subtype ‘forelimb wing’ (**Figure 5**). They do not expect historical homologs of the serial homolog (‘pectoral fin’) to be returned. Further, they do not expect phenotypes to parts of the homolog’s serial homolog (e.g., ‘humerus’ of the ‘forelimb’), or their developmental precursor (‘forelimb bud’ and ‘forelimb wing bud’).

**Figure 5.**
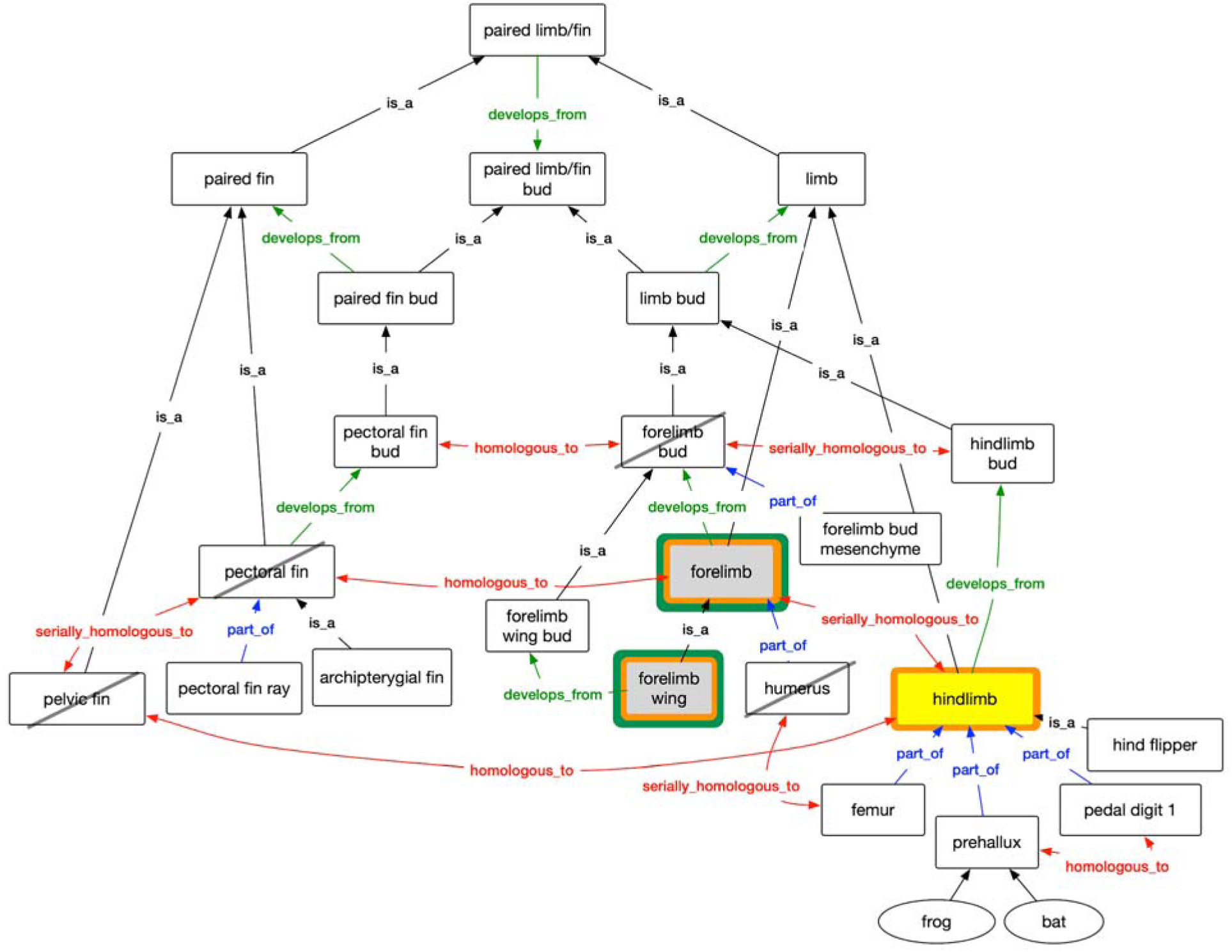
Competency question 5. Representation of pectoral and pelvic appendage structures in the Uberon anatomy ontology. Serial homology query for ‘hindlimb’ (bright yellow fill). Our persona expected ‘forelimb’ and ‘forelimb wing’ to be returned (grey fill). They did not expect parts of the serial homolog (‘humerus’), historical homologs of the serial homolog (‘pectoral fin’), or the developmental precursor of the serial homolog (‘forelimb bud’). Further, they did not expect the historical homolog of ‘hindlimb’, ‘pelvic fin’, to be returned in serial homology query. REA (green outline) and AVA (orange outline) returned the expected results, and AVA additionally returned ‘hindlimb’.

#### Competency question 6

Our persona expects that a query for serial homologs of ‘**hindlimb bud**’ would return phenotypes for ‘forelimb bud’ and its subtype ‘forelimb wing bud’ (**Figure 6**), but not its developmental product ‘forelimb’, nor its parts (‘forelimb bud mesenchyme’), or its serial homolog (‘pectoral fin bud’).

**Figure 6.**
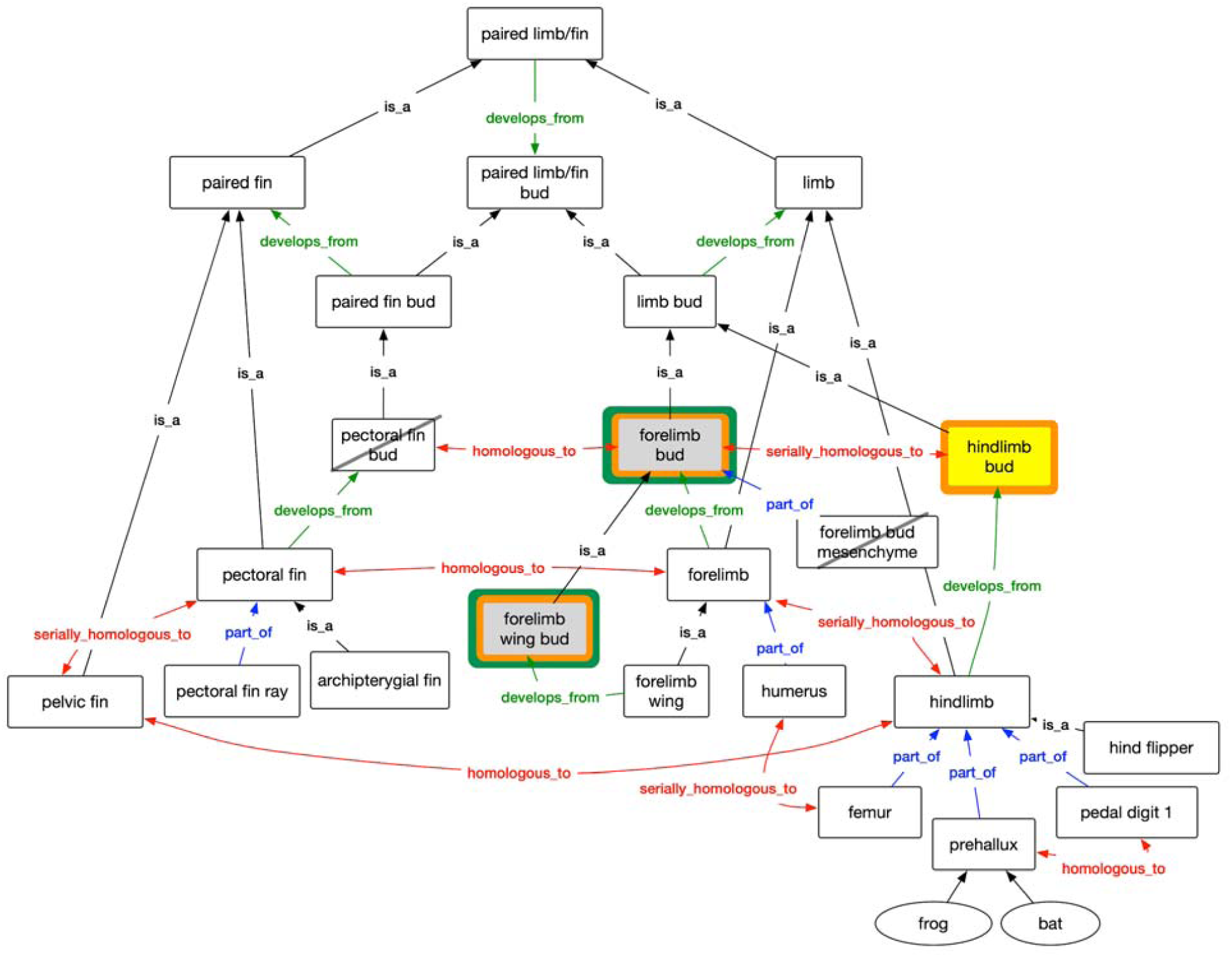
Competency question 6. Representation of pectoral and pelvic appendage structures in the Uberon anatomy ontology. Serial homology query for ‘hindlimb bud’ (bright yellow fill). Our persona expected ‘forelimb bud’ and ‘forelimb wing bud’ to be returned (grey fill). They did not expect (black slash) parts of the serial homolog (‘forelimb bud mesenchyme’) or historical homologs (‘pectoral fin bud’) of the serial homolog. REA (green outline) and AVA (orange outline) returned the expected results, and AVA additionally returned ‘hindlimb bud’.

#### Competency question 7

Our persona expects a phenotype query for serial homologs of **‘hind flipper’** to return phenotypes for ‘hindlimb’, its serial homolog ‘forelimb’, and subclasses of ‘forelimb’ such as ‘forelimb wing’, as illustrated in **Figure 7**. They do not expect the query to return structures for parts (e.g., ‘humerus’, ‘femur’) of the serial homologs, historical homologs (e.g., ‘pectoral fin’) of the serial homologs, or developmental precursors (e.g., ‘hindlimb bud’, ‘forelimb bud’) of the serial homologs.

**Figure 7.**
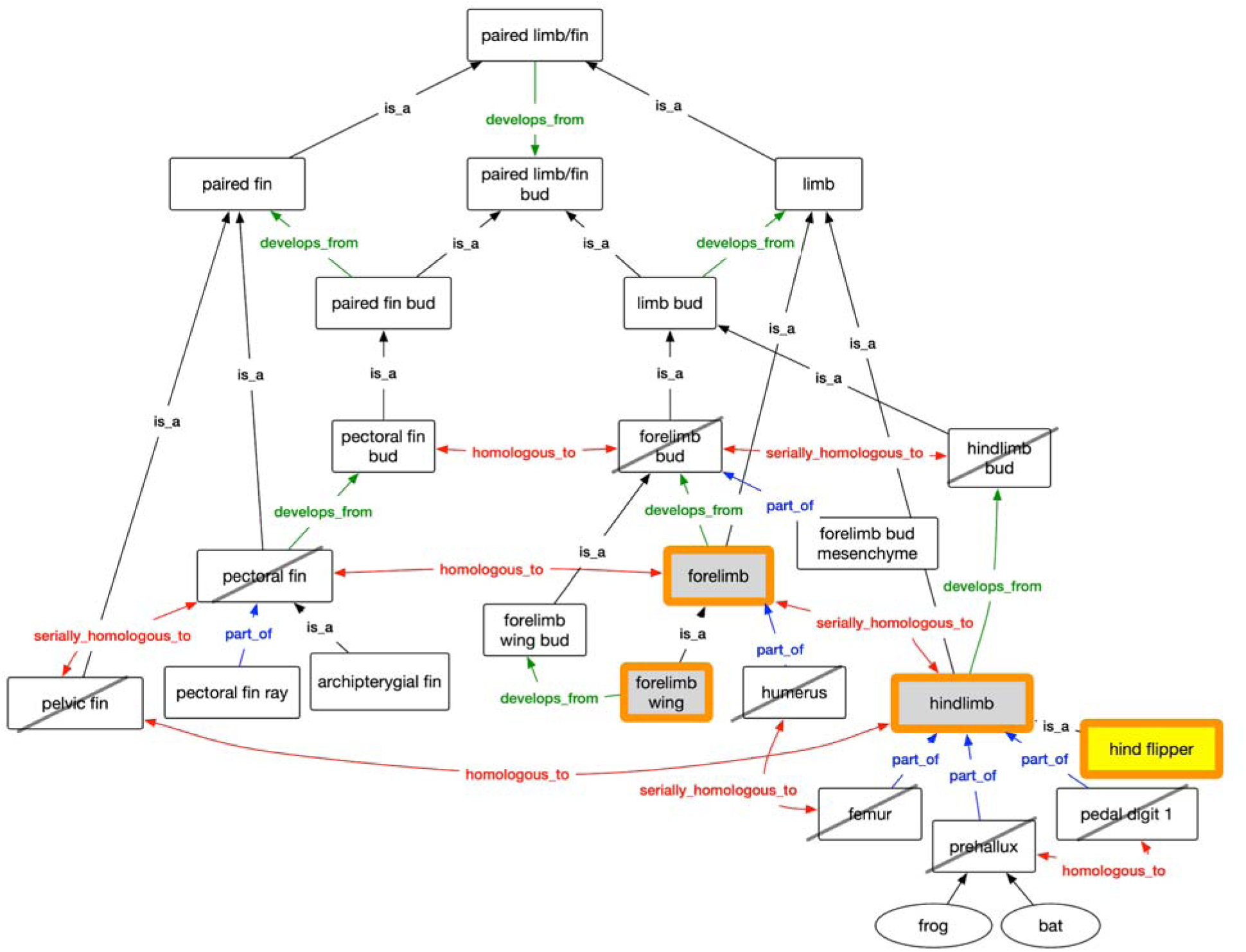
Competency question 7. Representation of pectoral and pelvic appendage structures in the Uberon anatomy ontology. Serial homology query for ‘hind flipper’ (bright yellow fill). Our persona expected ‘hindlimb’, ‘forelimb’, and ‘forelimb wing’ to be returned (grey fill). Structures not expected (black slash) include parts of the serial homologs (e.g., ‘femur’, ‘prehallux’, ‘pedal digit 1’, ‘humerus’), historical homologs of the serial homolog (‘pectoral fin’, ‘pelvic fin’), and developmental precursors of the serial homolog (‘forelimb bud’, ‘hindlimb bud’). The results of the AVA query (orange outline) included expected structures in addition to the search term. In REA, no results were returned.

### Annotation of homology assertions

Assertions of homology and statements of lack thereof among the skeletal elements of vertebrates were extracted from the phylogenetic literature on teleost fishes and early sarcopterygians [12], reviews of fin and limb evolution [51,52], and select papers from the developmental genetic literature (e.g., [53,54]). We systematically sought explicit homology statements between the skeletal elements in actinopterygian fins and sarcopterygian fins and limbs. Though not comprehensive, because the literature in the area of fin/limb evolution is substantial and homology statements specific to many taxonomic groups were not extracted (e.g., between urodele and anuran amphibians), many of the well-known and controversial homologies across fishes and amphibians were captured.

Similar to [55], we found that evidence for homology is explicitly asserted in the literature only rarely. This is particularly surprising in the phylogenetic literature, where what is judged to be the same character state represents an explicit hypothesis of putative primary homology among the taxa that share it. Although investigators routinely judge sameness (homology) using criteria of similarity in structure or topographic position, in relation to specific character states, this is rarely explicitly stated. That is, a statement such as ‘Anatomical feature X in taxa A, B, and C is similar in structure and they are thus considered homologous’ is rare in the literature. An additional issue was observed in extracting homology statements from the comparative monographic and fin to limb evolution review literature, in that the focus is often on skeletal elements where homologies are not clear (e.g., radials, digits) as compared to elements such as the humerus or femur, where the homologies are thought to be clear (though rarely explicitly described).

Homology statements were annotated using the appropriate ontologies: anatomy terms using the Uberon anatomy ontology [5] and taxa with the Vertebrate Taxonomy Ontology [6]. Along with attribution for each statement, we recorded the type of evidence that supported or rejected a historical or serial homology relationship [2] using the following terms from the Evidence and Conclusion Ontology [56]: position (ECO:0000060), composition (ECO:0000063), development (ECO:0000067), morphology (ECO:0000071), and gene expression (ECO:0000075). We also recorded statements of homology for which a source of the evidence was cited and for which no evidence or source was explicitly given by annotation with the terms ‘traceable author statement’ (ECO:0000033) and ‘non-traceable author statement’ (ECO:0000034), respectively.

Assertions about homology in the literature sometimes also take the form of rejecting or discounting a homology relationship between structures. We recorded these, including the supporting evidence types as per above, using ‘not homologous to’, rather than ‘homologous to’, in the relationship column. While it is possible to encode the negation of a homology relationship in OWL, we typically only have these “not” assertions when there is a corresponding ‘is homologous_to’ statement. Adding the ‘not’ annotations as logical axioms would cause reasoning contradictions; thus we store these annotations as metadata, such that they don’t participate in reasoning.

The full collection of homology assertions, including fin and limb assertions, is publicly available at http://purl.org/phenoscape/demo/phenoscape_homology.owl and in Supplementary Materials 1. The fin/limb-specific homology assertions are shown in **Table 2**.

**Table 2.** The subset of homology assertions used in the present study pertaining to fins, limbs, and related structures. Each assertion relates an Entity 1 in Taxon 1 as historically or serially homologous (or not) to an Entity 2 in Taxon 2 based on evidence (annotated with terms from the Evidence and Conclusion Ontology [56]) cited in the literature (Attribution).

| Entity 1 | Taxon 1 | Relationship | Entity 2 | Taxon 2 | Evidence | Attribution |
| --- | --- | --- | --- | --- | --- | --- |
| **humerus** | Tetrapoda | *Serially homologous to* | **femur** | Tetrapoda | Gene expression | [57] |
| **forelimb** | Tetrapoda | *Serially homologous to* | **hindlimb** | Tetrapoda | Position | [58] |
| **forelimb bud** | Tetrapoda | *Serially homologous to* | **hindlimb bud** | Tetrapoda | Gene expression | [51,59] |
| **pectoral fin** | Vertebrata | *Homologous to* | **forelimb** | Tetrapoda | NAS | [60] |
| **pectoral fin** | Vertebrata | *Serially homologous to* | **pelvic fin** | Vertebrata | Gene expression | [51,59] |
| **pectoral fin bud** | Vertebrata | *Homologous to* | **forelimb bud** | Tetrapoda | Developmental similarity | [51,61] |
| **pelvic fin bud** | Vertebrata | *Homologous to* | **hindlimb bud** | Tetrapoda | Developmental similarity | [51,61] |
| **pelvic fin** | Vertebrata | *Homologous to* | **hindlimb** | Tetrapoda | NAS | [60] |
| **prehallux** | Anura | *Homologous to* | **pedal digit 1** | Tetrapoda | Developmental similarity | [49] |
| **prehallux** | Anura | *Not homologous to* | **pedal digit 1** | Tetrapoda | Developmental similarity | [50] |
| **prehallux** | Anura | *Not homologous to* | **pedal digit 1** | Tetrapoda | Developmental similarity | [49] |

We did not include homology axioms from the Vertebrate Homologous Organ Group ontology (vHOG) [39], because most of these axioms (as of 23 July 2018) are ‘self-homologies’ with a specific taxonomic scope. For example, vHOG represents the humerus bone as a historically homologous structure within taxon Sarcopterygii. Because in OWL each class is also a subclass of itself, this approach does not yield any additional results (*i.e.*, logical entailments). For the purposes of our investigation, they are redundant with the axioms provided by the anatomy ontology.

### Homology demonstration file

To evaluate the two homology models and demonstrate how they differ, we assembled a set of phenotypes for fish fins and tetrapod limbs and a corresponding set of homology assertions among the relevant entities. Twenty ontology-annotated phenotypes for entities that are types of fin and limb and their literature sources included in the demonstration file were drawn from the >72,000 gene and taxon phenotypes in the Phenoscape KB. An additional three phenotypes (‘small forelimb buds’, ‘forelimb wing bud present’, ‘forelimb bud mesenchyme present’) were also added to the demonstration file for testing purposes. This set of 23 fin/limb phenotypes used in the demonstration file are shown in **Table 3** and in Supplementary Materials 2. OWL instances representing organism phenotype annotations were created using the Protégé OWL editor, following the Entity–Quality model [47,48]. For each competency question, we added a named class expression to this OWL file, for the purpose of allowing an automated reasoner to infer subsumption of phenotype instances. The homology demonstration file was provisioned with the expected phenotypes as well as phenotypes that would not be expected, since biologists may also have expectations of results that should not be returned (e.g., parts or developmental precursors).

**Table 3.**
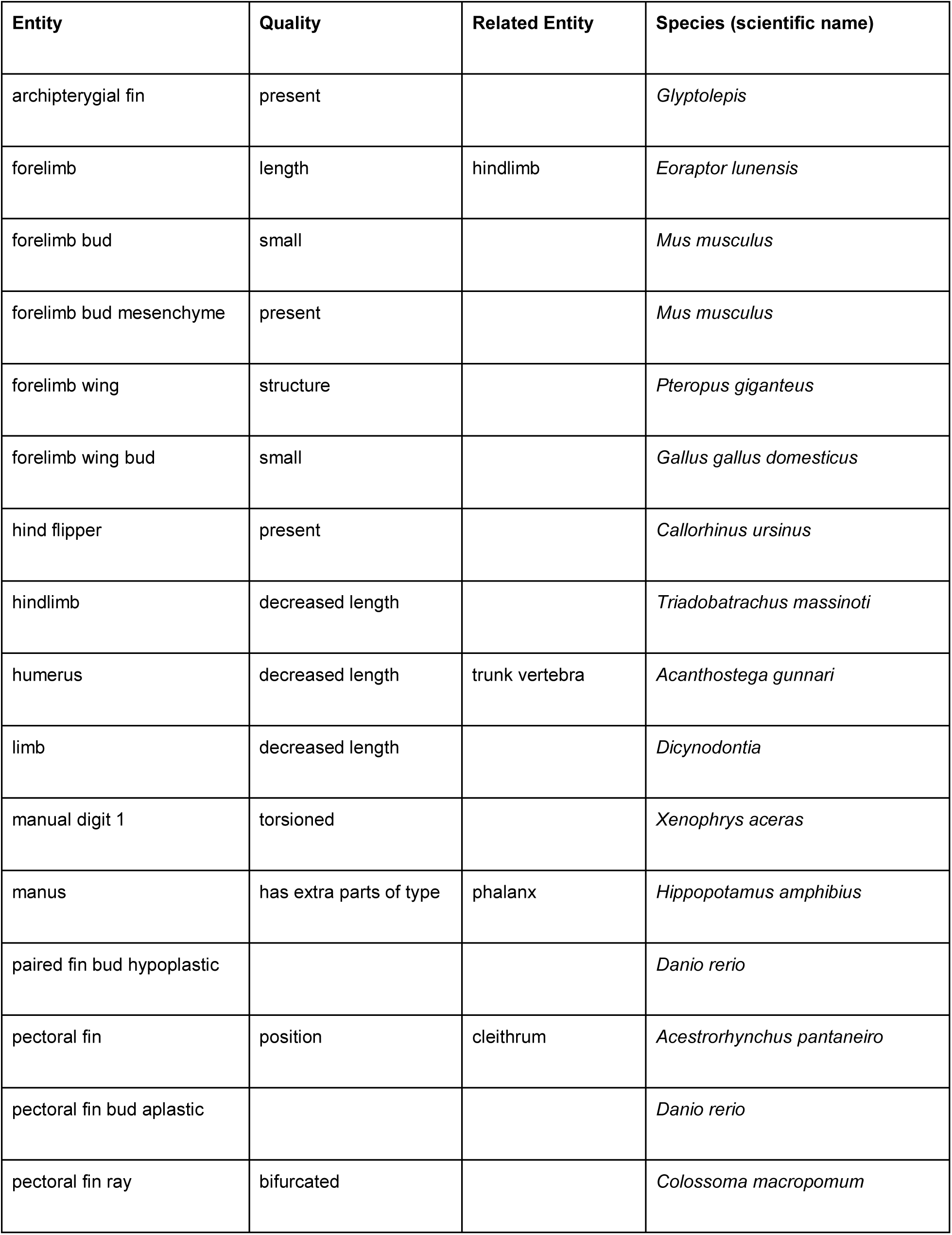
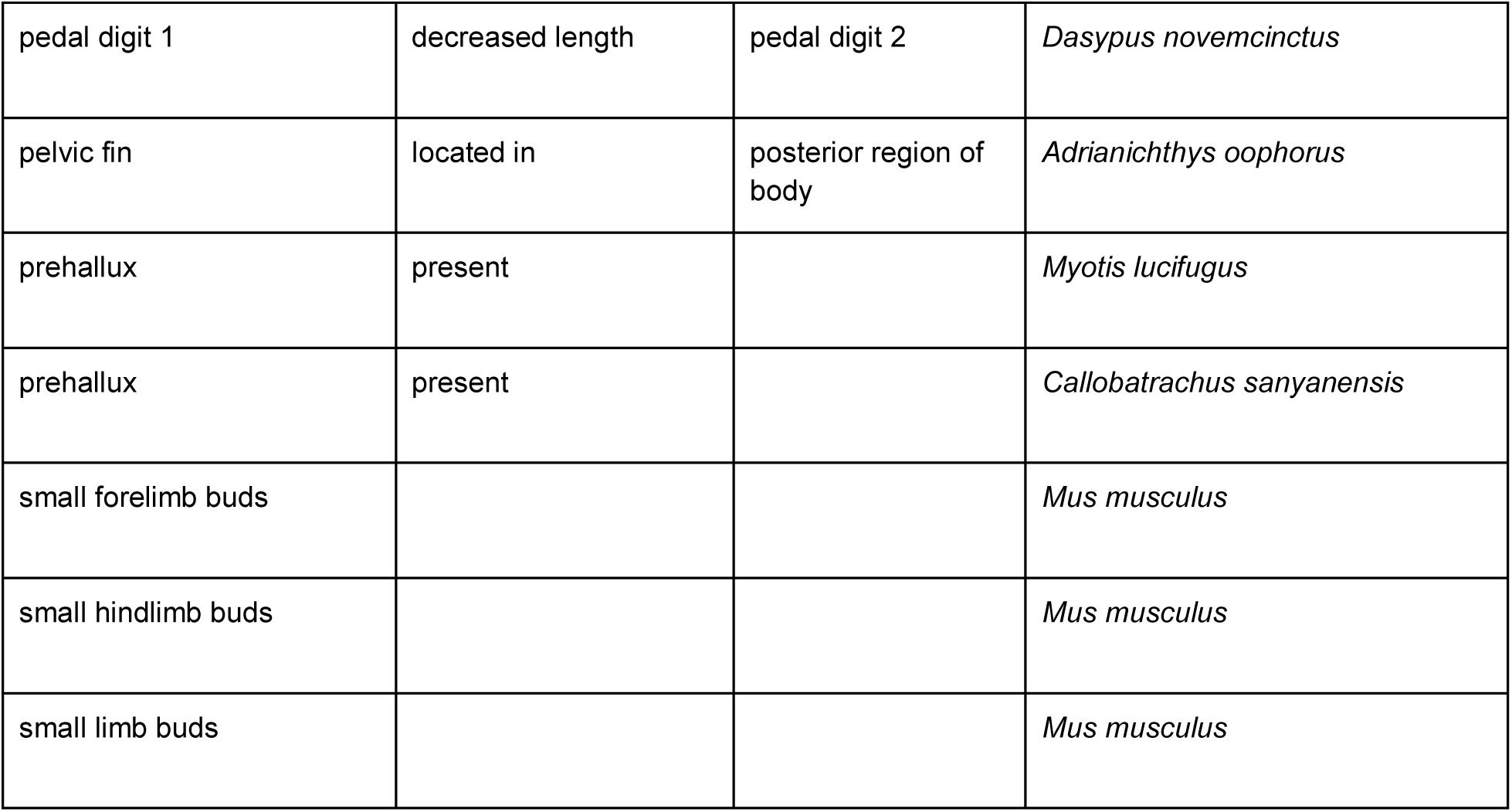
Subset of fin/limb phenotypes from the Phenoscape Knowledgebase (KB) used in the demonstration ontology to test the REA vs. AVA homology models.

We generated sets of OWL axioms representing homology relationships for each of the two different models (REA and AVA) using Scala scripts included in the phenoscape-owl-tools project (https://github.com/phenoscape/phenoscape-owl-tools). We used the ROBOT command-line tool (http://robot.obolibrary.org) to extract a reduced module of axioms from the Uberon anatomy ontology relevant to the terms used in the demonstration ontology, using syntactic locality module extraction, [62], and to construct a merged ontology file for each homology model. These merged files are small enough to be queried within Protégé using the HermiT OWL-DL reasoner [63], which comes with Protégé. The demonstration ontology workflow is available on GitHub in the homology-annotations-demo project (https://github.com/phenoscape/homology-annotations-demo).

## Results

### Homology statements

In total 46 homology assertions were collected for the paired fins and limbs, including ten statements pertaining to serial homology. Six positive assertions of homology were contradicted by negative statements of homology. For example, the alular digit in birds was asserted as homologous to manual digit 1 in non-avian tetrapods based on gene expression evidence [54,64], whereas these two structures were deemed not homologous based on developmental and morphological similarity [31]. The most common evidence type recorded was based on development (27 statements), followed by morphological similarity (26 statements), position (20 statements), and gene expression (14 homology statements); 5 statements cited evidence traceable to a different publication, whereas 6 statements did not cite traceable evidence.

### REA vs. AVA models

Results from REA and AVA models (**Figures 1-7**) are described in relation to each competency question and in **Table 4.**

**Table 4.** The results expected by our persona and the results obtained under REA and AVA models for competency questions one through seven. Results from AVA that are due to self-homology and subtype are denoted in italics.

| Competency question | Query term | Homology | Expectation | REA results | AVA results |
| --- | --- | --- | --- | --- | --- |
| 1 | Pectoral fin (Figure 1) | Historical | Forelimb<br>Forelimb wing | Forelimb<br>Forelimb wing | Forelimb<br>Forelimb wing<br>*Pectoral fin*<br>*Archipterygial fin* |
| 2 | Forelimb wing (Figure 2) | Historical | Forelimb<br>Pectoral fin<br>Archipterygial fin | none | Forelimb<br>Pectoral fin<br>Archipterygial fin<br>*Forelimb wing* |
| 3 | Pectoral fin bud (Figure 3) | Historical | Forelimb bud<br>Forelimb wing bud | Forelimb bud<br>Forelimb wing bud | Forelimb bud<br>Forelimb wing bud<br>*Pectoral fin bud* |
| 4 | Pedal digit 1 (Figure 4) | Historical | Prehallux in anurans (frogs) | Prehallux in anurans (frogs) | Prehallux in anurans (frogs)<br>*Pedal digit 1* |
| 5 | Hindlimb (Figure 5) | Serial | Forelimb<br>Forelimb wing | Forelimb<br>Forelimb wing | Forelimb<br>Forelimb wing<br>*Hindlimb*<br>*Hind flipper* |
| 6 | Hindlimb bud (Figure 6) | Serial | Forelimb bud<br>Forelimb wing bud | Forelimb bud<br>Forelimb wing bud | Forelimb bud<br>Forelimb wing bud<br>*Hindlimb bud* |
| 7 | Hind flipper (Figure 7) | Serial | Hindlimb<br>Forelimb<br>Forelimb wing | none | Hindlimb<br>Forelimb<br>Forelimb wing<br>*Hind flipper* |

### Competency question 1 results

Both REA and AVA returned the expected phenotypes; AVA additionally returned phenotypes for the search term itself (‘pectoral fin’) and its subtype (‘archipterygial fin’) (**Figure 1**) (**Table 4**). These results are consistent with OWL entailments of the respective models.

### Competency question 2 results

REA did not return any results from a query for homologs of ‘forelimb wing’. The same AVA query returned all expected results and additionally phenotypes for the search term itself (‘forelimb wing’) (**Figure 2**) (**Table 4**). These results are consistent with OWL entailments of the respective models.

### Competency question 3 results

Both REA and AVA returned ‘forelimb bud’ and ‘forelimb wing bud’; AVA returned the expected results and additionally ‘pectoral fin bud’ (**Figure 3**) (**Table 4**). These results are consistent with OWL entailments of the respective models.

### Competency question 4 results

REA returned the expected results; AVA returned the expected results and additionally ‘pedal digit 1’ (**Figure 4**) (**Table 4**). These results are consistent with OWL entailments of the respective models.

### Competency question 5 results

REA returned the expected results. AVA returned the expected results and additionally ‘hindlimb’ (**Figure 5**) (**Table 4**). These results are consistent with OWL entailments of the respective models.

### Competency question 6 results

REA returned the expected results; AVA returned the expected results and additionally ‘hindlimb bud’ (**Figure 6**) (**Table 4**). These results are consistent with OWL entailments of the respective models.

### Competency question 7 results

REA returned no results from a query for homologs of ‘hind flipper’. The same AVA query returned all expected results and additionally phenotypes for the search term itself (‘hind flipper’) (**Figure 7**) (**Table 4**). These results are consistent with OWL entailments of the respective models.

## Discussion

Effectively incorporating homology relationships into anatomy ontologies lays the groundwork for this knowledge to be used in other ontology-based tools and reasoning applications, including candidate gene discovery and phenotypic matrix assembly. We developed and evaluated two models for representing historical and serial homology relations using a collection of homology assertions and a set of taxon phenotypes for vertebrate fins and limbs from the Phenoscape Knowledgebase (KB). The two models that we evaluated reflect an inherent tradeoff between expressivity and computational efficiency. While there are other ways to represent homology, these two models are sufficient to show that there are logical ramifications that differ in ways that might surprise many biologists. We have surfaced these differences by way of competency questions that force us to specify exactly what a biologist would expect by way of reasoning outcome. These expectations may differ among biologists, and our competency questions are not comprehensive, but we believe that we have provided a foundation that can be built upon by future investigators.

Both of the OWL models we explored represented homology assertions as a binary relation. For example, we represented the homology statement as ‘forelimb wings in birds are homologous to pectoral fins in fishes’. Homology can also be considered as a ternary relation [15] which points the two homologs (*e.g.*, forelimb wings, pectoral fins) to a more general reference point -- the ancestral structure from which they evolved (in this case ‘pectoral appendages’) in the named monophyletic group that encompasses them. Bock [15] argues that this conditional phase describing the condition of the feature in the common ancestor should always be included in any statement about the homology of features. For example, the wings of birds and the wings of bats are homologous as tetrapod forelimbs -- or the wings in birds are homologous to the pectoral fins in fishes as vertebrate pectoral appendages. In practice it is often difficult to conceptualize and describe an ancestral anatomical structure in detail; the only description possible is often only at a high level. For example, the hyomandibula in the jaw of fishes is homologous to the stapes, an inner ear bone in mammals, as a bone of the dorsal hyoid arch in vertebrates. ‘Bone of the dorsal hyoid arch’, references some bone in a region and is not more informative than an ontological parent class expression such as ‘endochondral bone that is part of or derived from the hyoid arch skeleton’. That is, in the binary representation, the homologs are also connected to a more general anatomical class, but there it is implied by the structure of the ontology and is thus not necessarily an evolutionary concept. Further practical difficulties with ternary representation arise in pointing to the common ancestor from which both homologous structures arose. It may not be possible to determine the position of this ancestor if there is incongruence among phylogenetic trees. Even where there is a single robust phylogeny, there may not be a named taxon class or other identifier that corresponds to the last common ancestor. The binary representation does not point to the last common ancestor for the taxa bearing the homologous anatomical structures (Tetrapoda or Vertebrata in the above examples). However, because anatomical annotations are to taxa, the data can be referred to a phylogeny of choice to infer the ancestral taxon. An additional reason for representing homology statements using binary relations is because ternary relations are more complicated and awkward to use and query in OWL. However, if desirable and practical in the future, the AVA model could support the ternary representation by specifying an ontological class of the ancestral structure.

### Evaluating homology models

Both the REA and AVA models returned the user-expected results for all but one historical homology query and all serial homology queries. For each competency question, the AVA model also returned the specified query term and any subtypes, because the query term is itself a descendant of the ancestral structure in the model. That the expected results were not returned for Competency Question 2 (query for homologs of ‘forelimb wing’, Figure 2) is a result of our choice to model homology using existential property restrictions of the REA model. Specifically, the relevant homology axiom for this query states that every ‘pectoral fin’ is homologous_to *some* ‘forelimb’. It cannot be assumed by an OWL reasoner that the ‘forelimb’ being referred to is a ‘forelimb wing’; it may be some other subtype of ‘forelimb’. Thus, under the REA model, no results were returned from this query. In the AVA model, however, the semantics are defined such that “every ‘pectoral fin’ is homologous_to *every* ‘forelimb’”, and thus ‘forelimb wing’ was returned.

Although the AVA model more closely meets our persona’s expectations for the competency questions, its reliance on more expressive OWL reasoning prohibits its use in practice, e.g., at the scale of a knowledgebase such as the Phenoscape KB. As discussed above in “Logical models of homology assertions”, the REA model is amenable to more efficient reasoning, such as with the ELK reasoner. Furthermore, although some of our persona’s expected results were missing, REA returned no incorrect answers for our test data.

### Formalizing homology

Although considerable research and thought has been applied to understanding how homology can be identified and further codified, general expectations for a semantic model have not been previously formalized. We translated this biological knowledge into the framework of an ontology graph, considering carefully the way in which homology relationships would be expected to propagate along the logical relationships among entities, their subtypes, parts, and developmental precursors and products.

For example, although in some cases the parts of homologous structures might be homologous, they often are not, and thus homology is not propagated through parthood relationships in our models. This is the case even for serial homologs. No biologist has generally surmised, for instance, that skeletal parts of the fish pectoral fins are homologous to those of our forelimbs, though some have suggested homology between specific parts (radials of the fin to humerus of the forelimb). Incorrect inferences are therefore not realized in our semantic model. Rather, where applicable, historical or serial homology must be directly asserted between structures that are parts of homologs. For example, ‘humerus’ (part of the forelimb) and ‘radial’ (part of the pectoral fin) need to be directly asserted as homologs, even if the structures of which they are a part, ‘pectoral fin’ and ‘forelimb’, are already asserted as homologs.

Another example of limiting homology inference on the basis of biological knowledge comes from developmental biology. Here we restricted reasoning across development because of the widely recognized disconnect between homology at different levels of biological organization: homology at one level does not necessitate homology at another [65–67]. There are many examples of homologous structures that develop from non-homologous developmental precursors [18]. For example, Meckel’s cartilage (part of the jaw) in vertebrates is induced differently in amphibians, birds, and mammals [18]. *Vice versa*, there many examples of non-homologous structures whose development is similar, e.g., under the control of orthologous genes. For example *distal-less* regulates outgrowth of the limbs of insects and vertebrates, but phylogenies nearly conclusively reflect the independent evolution of limbs in these taxa (i.e., they are not historical homologs) [68]. Because of this lack of homology correspondence across biological levels, the desired outcome from a query for historical homologs of ‘pectoral fin bud’ would be ‘forelimb bud’ or ‘forelimb wing bud’, but not the product of further bud development, *i.e*., ‘forelimb’ or ‘forelimb wing’. *Vice versa*, the desired outcome from a query for historical homologs of ‘pectoral fin’ would be ‘forelimb’ and its subtype ‘forelimb wing’, but not their developmental precursors ‘forelimb bud’ and ‘forelimb wing bud’.

We also took a conservative approach to extending reasoning across multiple types of homology relationships. For example, although ‘hindlimb’ is serially homologous to ‘forelimb’ and ‘hindlimb’ is historically homologous to ‘pelvic fin’, a query for serial homologs of ‘hindlimb’, returned its serial homolog ‘forelimb’, but not the historical homolog of forelimb, *i.e.*, ‘pectoral fin’. Thus, a serial homology search does not extend to historical homologs of the serial homolog, and likewise an historical homology search does not extend to serial homologs of the historical homolog.

### Homology assertions must be specific

In initial tests of the reasoning based on homology assertions from the literature we discovered that homology axioms involving general, *i.e.*, less specific, grouping terms can return unexpected results. For example, although it is accepted that the paired fins of fishes are homologous to the limbs of terrestrial vertebrates, when this statement is translated into a homology assertion (‘paired fin’ homologous to ‘limb’), the queries involving the more specific subtypes of these terms yield some erroneous results. Under the AVA model, a query for homologs of ‘pectoral fin’ return both ‘forelimb’ and ‘hindlimb’ because of the semantics of the homology axiom: *every* ‘paired fin’ homologous to *every* ‘limb’. Here, because pectoral fin is a type of paired fin, and under the relationship where paired fin is homologous to limb, the outcome includes both subtypes of ‘limb’, the forelimb (true historical homolog) and hindlimb (not historical homolog). In contrast, the REA model only returned ‘forelimb’ (and subtype ‘forelimb wing’), because the semantics for this model (*every* ‘paired fin’ homologous to *some* ‘limb’) asserts that only some instances of ‘limb’ are homologous to ‘paired fin’. Because pectoral fin also has a homology assertion to forelimb, only forelimb is returned.

Although in many cases it may suffice for a user to query for serial homologs by using a shared parent term (*e.g.*, a query for ‘vertebra’ returns ‘vertebra 1’, ‘vertebra 2’, ‘vertebra 3’, etc…), in other cases, explicit homology axioms are needed to relate serial homologs. For example, ‘humerus’ and ‘femur’ need an explicit homology axiom because these terms do not share a common parent term in Uberon. Other types of iterative homology [16], i.e., between bilaterally (*e.g.*, vertebrates) or radially symmetric (*e.g.*, echinoderms) structures or male vs. female organisms, also require explicit serial homology axioms. For example, although terms for structures between between right and left sides of the body are subtypes of the more general structure (*e.g.*, ‘right preopercle’ and ‘left preopercle’ are subtypes of ‘preopercle’), homology between them needs to be asserted. Without such specification, searches for these types of iterative homologs fail (*e.g.*, a search for homolog of the ‘right preopercle’ does not return the expected result, i.e., ‘left preopercle’).

### Homology grouping classes

The Uberon anatomy ontology contains ten explicit ‘grouping classes’ primarily driven by homology (as opposed to structure, function or position) [5]. These are high level classes of ‘nearly certain’ homology that were historically developed for Uberon to ensure that users received expected results from data queries without having to explicitly include homology assertions and a model that implements them. For example, a user query to ‘paired limb/fin’ would return ‘paired fin’ and ‘limb’ (**Figure 1**). These grouping classes are designated with the ‘in_subset: homology_grouping’ tag, but are not logically related as homologous and do not include evidence or attribution. Nine of these ten classes are relevant to the fin/limb collection of homology assertions assembled here: ‘paired limb/fin bud’ UBERON:000435; ‘limb/fin segment’ UBERON:0010538; ‘paired limb/fin cartilage’ UBERON:0007389; ‘paired limb/fin skeleton’ UBERON:0011582; ‘pelvic appendage’ UBERON:000470; ‘paired limb/fin’ UBERON:0004708; ‘pectoral appendage’ UBERON:0004710; ‘paired limb/fin field’ UBERON:0005732, and ‘bone of free limb or fin’ UBERON:0004375. These grouping classes do not affect the outcome of the reasoning (see Supplementary Materials 3).

### Disabling anatomical homology relations to discover deep homology

The discovery of similar anatomical features that arose independently in evolution and yet are underlain by homologous genes and networks has given pause to many investigators focused on homology at the structural level. Such highly conserved genetic regulation, termed ‘deep homology’ [69,70] reflects not only the deep continuity of fundamental circuitry across long stretches of evolution, but also its co-option to generate similar anatomical structures that are non-homologous. The extent of deep homology across life is unclear, and it will be necessary to make many comparisons of similar structures across diverse organisms to gauge if it is the rule or the exception. Such a research program would be enhanced by the ability to conduct taxonomically broad similarity searches in a knowledge base such as the one used here. The results of interest, in this case, would be structures that do *not* owe their similarity to historical or serial homology, such as fly wings, vertebrate limbs and beetle horns as ‘appendages’, or the light sensing organs of arthropods, molluscs and vertebrates as ‘eyes’. Thus, implementation of the homology axioms described herein may be useful in providing either a negative or positive filter for search results, depending on the application.

### Implementation in the Phenoscape KB

We have incorporated historical and serial homology reasoning in the Phenoscape KB, where it allows discovery of structures that are related because of common ancestry (**Figure 8**). Fully implementing homology queries, however, still remains a challenge owing to the limitations of OWL reasoning. In the Phenoscape KB, for example, the more computationally feasible REA model of homology was implemented. However, given the size of the anatomy and phenotype ontologies used by Phenoscape, even with REA, OWL reasoning on the complete terminology is only feasible using fast EL reasoners such as ELK [71]. Although we ultimately select and deploy a model that satisfies basic reasoning, we expect that it can and will be optimized for different purposes and as computational methods to represent uncertainty, hierarchical trait dependencies, and other variables evolve [72].

**Figure 8.**
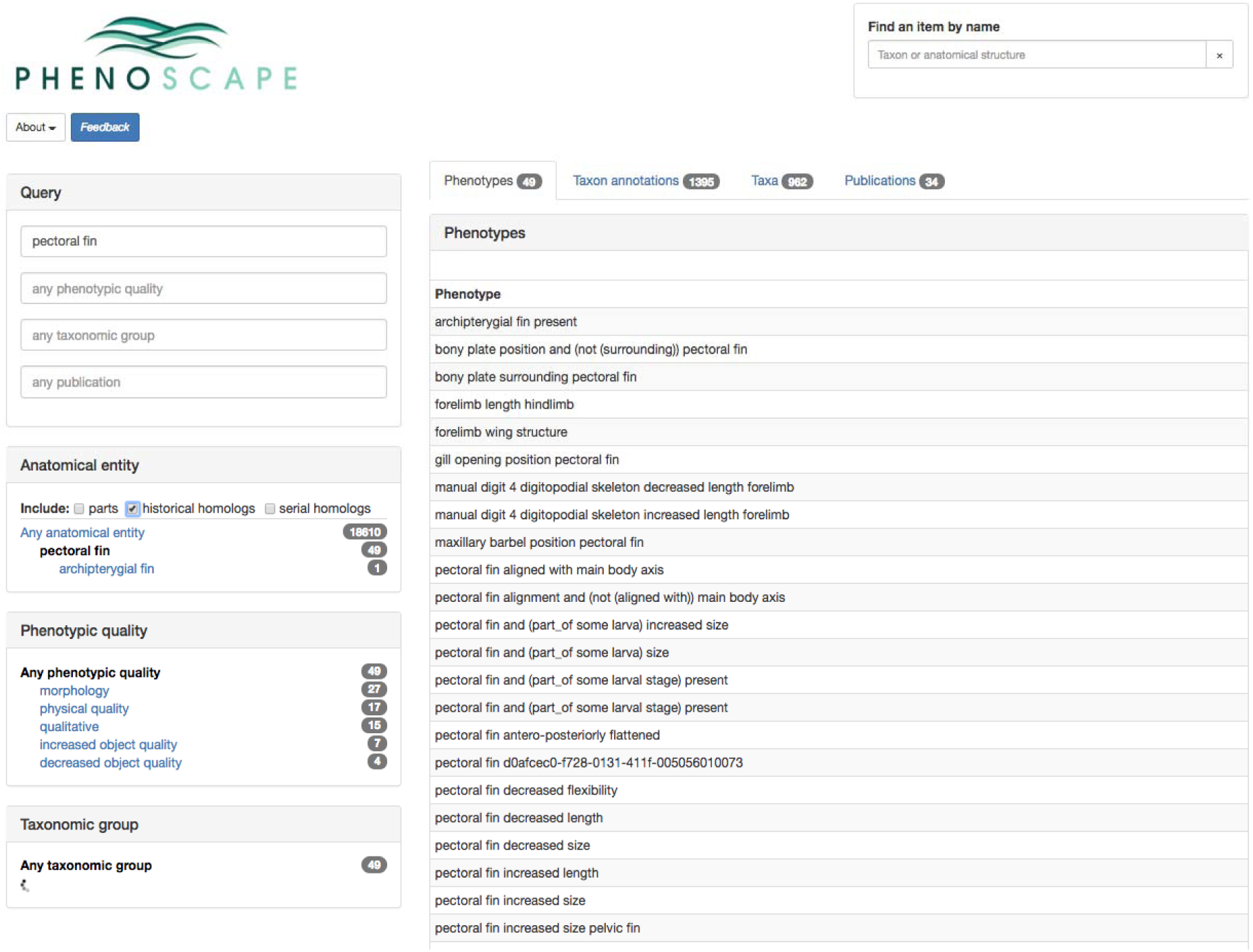
The results of historical homology reasoning in the Phenoscape Knowledgebase. A query for ‘pectoral fin’ phenotypes returns annotations to the distinct homolog class ‘forelimb’ as well as annotations to the original class (‘pectoral fin’) and its subtypes (in this case, ‘archipterygial fin’).

In the Phenoscape KB, user queries are currently restricted to positive homology assertions for both historical and serial homologs because contradictory statements cannot be used in reasoning. We previously envisioned enabling a user to choose the specific set of homology assertions to inform their searches [2]. Translating this into a functional model, however, is challenging because of the difficulty of representing conflicting statements (*i.e*. both homologous_to and not_homologous_to for the same pair of anatomical entities) within ontologies. Part of this challenge is that reasoning required to handle ‘not homologous to’ annotations is not implemented in the Phenoscape homology model. This is because these negative homology assertions are nearly always paired with disagreeing positive homology assertions. Including contradictory assertions in the logical definition of a single anatomical class would render that class unsatisfiable and thus unusable for data retrieval. However, ‘not homologous to’ relations are displayed in the metadata in the KB for anatomical terms. Although representing both positive and negative assertions might potentially serve the avid and discriminating comparative anatomist, enabling such choices would not necessarily be relevant to users from many other backgrounds.

### Modifying homology assumptions on-the-fly

Consensus concerning the homology of many structures may never be achieved, as different lines of evidence can point in opposing directions. As described above, whether the first ‘finger’ of birds is homologous or not to that in dinosaurs is a well known example of conflicting evidence. Although we relate homology assertions herein to the data that support them by annotation with homology evidence codes [2] from the Evidence & Conclusion Ontology (ECO) [73,74], they are not implemented in Phenoscape for customized homology reasoning. It may be desirable in the future, however, to allow user selection of homology assertions based on these codes. For instance, only homology relationships that are backed up by particular lines of evidence (*e.g*., ‘similarity of development’) might be chosen, or perhaps only homology that is supported by all the evidence (no conflict). Enabling individualized selection of homology relationships would alter the reasoning and thus the derivative products of knowledgebases. One would expect different sets of phenotypes to be reasoned, *e.g.*, using OntoTrace [12], and aggregated based on how similarity is treated, i.e., whether judged homologous or not. In turn, products derived from these phenotypes will be affected. These include hypotheses of evolution (phylogenies), candidates for genetic control elements [75] and genes [10], or even phenotype-based genomic identification [76] for biodiverse species. Because of the iterative nature of homology hypothesis development, these products may provide new evidence for the common ancestry vs. convergent nature of particular features. In summary, enabling machines to reason the various types of similarity (evolutionary, structural, functional, etc.) is a challenging but promising area for future work in the area of phenotype-driven knowledge discovery.

## Acknowledgments

The Phenoscape project was incubated starting in 2007 at the National Evolutionary Synthesis Center (NSF 0905606 and 0423641) and has been directly supported by NSF awards 1062404, 1062542, 0641025, 1661529. Phenoscape has been made possible by many collaborators and contributors (see full list at https://wiki.phenoscape.org/wiki/Acknowledgments). In particular, we thank M. Westerfield, M. Coburn, M.A. Haendel, J.G. Lundberg, and R. Mayden for discussions of homology and anatomy ontologies. We also thank the Phenotype Ontology Research Coordination Network (NSF 0956049) for support of a 2012 vertebrate working group meeting on homology that informed the present work. Discussion among the authors and D. Blackburn, T.A. Dececchi, H. Larsson, and K. Sears contributed many use cases that influenced our understanding of how homology should be represented. This manuscript is based in part on work done by P.M.M. while serving at the U.S. National Science Foundation. The views expressed in this paper do not necessarily reflect those of the National Science Foundation or the United States Government.

## Software and data availability

A snapshot of the code available at https://github.com/phenoscape/phenoscape-owl-tools and used in this paper is available from Zenodo: https://doi.org/10.5281/zenodo.2605467. The version of the demonstration ontology workflow available at https://github.com/phenoscape/homology-annotations-demo and used in this paper is archived at Zenodo (https://doi.org/10.5281/zenodo.2605488). These files can be run in Protégé with the HermiT reasoner [63]. The collection of Phenoscape homology assertions is available in Supplementary Materials 1, and the subset of fin/limb phenotypes used in the demonstration homology file is available in Supplementary Materials 2.

## Footnotes

1 Based on developmental comparisons with fossil early tetrapods, including *Ichthyostega* and *Eryops*, at least one study [49] considered the **prehallux**, i.e., the ossification anterior to the ‘big toe’ in frogs (anurans), to be a *bona fide* **digit**. That is, the homolog of the **prehallux** in anurans is **pedal digit 1** in *Ichthyostega* and *Eryops*. In the homology assertions file (Supplementary Materials 1) this is represented as “prehallux in Anura homologous to pedal digit 1 in Tetrapoda”, with the developmental similarity evidence code (ECO:0000067) and attribution to [49]. Other authors, e.g., [50] present developmental and variational evidence indicating that the prehallux is not the homolog of a digit, but rather a ‘predigit’. In the homology assertions file this is represented as “prehallux not homologous to pedal digit 1”, with the developmental similarity evidence code (ECO:0000067) and attribution to [50]. Further, a structure named ‘prehallux’ exists in non-anuran taxa, e.g., mammals, and the homology relationship across taxa is uncertain.

## References

1. Vogt L. Towards a semantic approach to numerical tree inference in phylogenetics. Cladistics. 2018;34: 200–224. doi:10.1111/cla.12195

2. Dahdul WM, Lundberg JG, Midford PE, Balhoff JP, Lapp H, Vision TJ, et al. The Teleost Anatomy Ontology: anatomical representation for the genomics age. Systematic Biology. 2010;59: 369–383. doi:10.1093/sysbio/syq013

3. Dahdul WM, Balhoff JP, Blackburn DC, Diehl AD, Haendel MA, Hall BK, et al. A unified anatomy ontology of the vertebrate skeletal system. PLoS One. 2012;7: e51070. doi:10.1371/journal.pone.0051070

4. Mungall CJ, Torniai C, Gkoutos GV, Lewis SE, Haendel MA. Uberon, an integrative multi-species anatomy ontology. Genome Biology. 2012;13: R5. doi:10.1186/gb-2012-13-1-r5

5. Haendel MA, Balhoff JP, Bastian FB, Blackburn DC, Blake JA, Bradford Y, et al. Unification of multi-species vertebrate anatomy ontologies for comparative biology in Uberon. Journal of Biomedical Semantics. 2014;5: 21. doi:10.1186/2041-1480-5-21

6. Midford PE, Dececchi TA, Balhoff JP, Dahdul WM, Ibrahim N, Lapp H, et al. The vertebrate taxonomy ontology: a framework for reasoning across model organism and species phenotypes. Journal of Biomedical Semantics. 2013;4: 34. doi:10.1186/2041-1480-4-34

7. Balhoff JP, Dahdul WM, Dececchi TA, Lapp H, Mabee PM, Vision TJ. Annotation of phenotypic diversity: decoupling data curation and ontology curation using Phenex. Journal of Biomedical Semantics. 2014;5: 45. doi:10.1186/2041-1480-5-45

8. Balhoff JP, Dahdul WM, Kothari CR, Lapp H, Lundberg JG, Mabee P, et al. Phenex: ontological annotation of phenotypic diversity. PLoS One. 2010;5: e10500. doi:10.1371/journal.pone.0010500

9. Mabee PM, Dahdul WM, Balhoff JP, Lapp H, Manda P, Uyeda J, et al. Phenoscape: Semantic analysis of organismal traits and genes yields insights in evolutionary biology. In: Thessen A, editor. Application of Semantic Technology in Biodiversity Science. Berlin: IOS Press; 2018.

10. Edmunds RC, Su B, Balhoff JP, Eames BF, Dahdul WM, Lapp H, et al. Phenoscape: identifying candidate genes for evolutionary phenotypes. Molecular Biology and Evolution. 2016;33: 13–24. doi:10.1093/molbev/msv223

11. Manda P, Mungall CJ, Balhoff JP, Lapp H, Vision TJ. Investigating the importance of anatomical homology for cross-species phenotype comparisons using semantic similarity. Proceedings of the Pacific Symposium on Biocomputing 2016. 2016. pp. 132–143. doi:10.1142/9789814749411_0013

12. Dececchi TA, Balhoff JP, Lapp H, Mabee PM. Toward synthesizing our knowledge of morphology: using ontologies and machine reasoning to extract presence/absence evolutionary phenotypes across studies. Systematic Biology. 2015;64: 936–952. doi:10.1093/sysbio/syv031

13. Jackson LM, Fernando PC, Hanscom JS, Balhoff JP, Mabee PM. Automated integration of trees and traits: a case study using paired fin loss across teleost fishes. Systematic Biology. 2018;67: 559–575. doi:10.1093/sysbio/syx098

14. Hall BK. Homology: The Hierarchical Basis of Comparative Biology. Academic Press; 2012.

15. Bock WJ. Discussion: the concept of homology. Annals of the New York Academy of Sciences. 1969;167: 71–73. doi:10.1111/j.1749-6632.1969.tb20434.x

16. Roth VL. On homology. Biological Journal of the Linnean Society. 1984;22: 13–29. doi:10.1111/j.1095-8312.1984.tb00796.x

17. Roth VL. The biological basis of homology. In: Humphries CJ, editor. Ontogeny and Systematics. New York: Columbia University Press; 1988. pp. 1–26.

18. Wagner GP. The biological homology concept. Annual Review of Ecology and Systematics. Annual Reviews; 1989;20: 51–69. doi:10.1146/annurev.es.20.110189.000411

19. Hall BK. Descent with modification: the unity underlying homology and homoplasy as seen through an analysis of development and evolution. Biological Reviews of the Cambridge Philosophical Society. 2003;78: 409–433. doi:10.1017/S1464793102006097

20. Minelli A, Fusco G. Homology. In: Kampourakis K, editor. The Philosophy of Biology History, Philosophy and Theory of the Life Sciences, vol 1. Dordrecht: Springer; 2013. pp. 289–322. doi:10.1007/978-94-007-6537-5_15

21. Scotland RW. Deep homology: a view from systematics. Bioessays. 2010;32: 438–449. doi:10.1002/bies.200900175

22. Kratochwil CF, Liang Y, Gerwin J, Woltering JM, Urban S, Henning F, et al. Agouti-related peptide 2 facilitates convergent evolution of stripe patterns across cichlid fish radiations. Science. 2018;362: 457–460. doi:10.1126/science.aao6809

23. Briscoe SD, Ragsdale CW. Homology, neocortex, and the evolution of developmental mechanisms. Science. 2018;362: 190–193. doi:10.1126/science.aau3711

24. Cracraft J. Phylogeny and evo-devo: characters, homology, and the historical analysis of the evolution of development. Zoology. 2005;108: 345–356. doi:10.1016/j.zool.2005.09.003

25. Roux J, Rosikiewicz M, Robinson-Rechavi M. What to compare and how: comparative transcriptomics for Evo-Devo. Journal of Experimental Zoology Part B-Molecular and Developmental. Wiley Online Library; 2015;324B: 372–382. doi:10.1002/jez.b.22618

26. Romer AS. The Vertebrate Body. Philadelphia: WB Saunders Company; 1950. pp. 888–888.

27. Mabee PM, Ashburner M, Cronk Q, Gkoutos GV, Haendel M, Segerdell E, et al. Phenotype ontologies: the bridge between genomics and evolution. Trends in Ecology & Evolution. 2007;22: 345–350. doi:10.1016/j.tree.2007.03.013

28. Mabee PM, Arratia G, Coburn M, Haendel M, Hilton EJ, Lundberg JG, et al. Connecting evolutionary morphology to genomics using ontologies: a case study from Cypriniformes including zebrafish. Journal of Experimental Zoology Part B-Molecular and Developmental Evolution. 2007;308B: 655–668. doi:10.1002/jez.b.21181

29. Mabee PM. Supraneural and predorsal bones in fishes: development and homologies. Copeia. 1988;1988: 827–838. doi:10.2307/1445705

30. Wagner GP, Gauthier JA. 1,2,3 = 2,3,4: a solution to the problem of the homology of the digits in the avian hand. Proceedings of the National Academy of Sciences. 1999;96: 5111–5116. doi:10.1073/pnas.96.9.5111

31. Burke AC, Feduccia A. Developmental patterns and the identification of homologies in the avian hand. Science. 1997;278: 666–668. doi:10.1126/science.278.5338.666

32. Feduccia A. 1,2,3 = 2,3,4: accommodating the cladogram. Proceedings of the National Academy of Sciences. 1999;96: 4740–4742. doi:10.1073/pnas.96.9.4740

33. Haendel MA, Neuhaus F, Osumi-Sutherland D, Mabee PM, Mejino JLV Jr, Mungall CJ, et al. CARO–the Common Anatomy Reference Ontology. In: Burger A, Davidson D, Baldock R, editors. Anatomy Ontologies for Bioinformatics. London: Springer; 2008. Available: https://doi.org/10.1007/978-1-84628-885-2_16

34. Travillian R, Malone J, Pang C, Hancock J, Holland PWH, Schofield P, et al. The Vertebrate Bridging Ontology (VBO). Bio-Ontologies 2011. 2011; Available: http://bio-ontologies.knowledgeblog.org/231

35. Malone J, Holloway E, Adamusiak T, Kapushesky M, Zheng J, Kolesnikov N, et al. Modeling sample variables with an Experimental Factor Ontology. Bioinformatics. 2010;26: 1112–1118. doi:10.1093/bioinformatics/btq099

36. Parmentier G, Bastian FB, Robinson-Rechavi M. Homolonto: generating homology relationships by pairwise alignment of ontologies and application to vertebrate anatomy. Bioinformatics. 2010;26: 1766–1771. doi:10.1093/bioinformatics/btq283

37. Bastian F, Parmentier G, Roux J, Moretti S, Laudet V, Robinson-Rechavi M. Bgee: Integrating and comparing heterogeneous transcriptome data among species. Data Integration in the Life Sciences DILS 2008 Lecture Notes in Computer Science, vol 5109. Springer, Berlin, Heidelberg; 2008. pp. 124–131. doi:10.1007/978-3-540-69828-9_12

38. Roux J, Robinson-Rechavi M. An ontology to clarify homology-related concepts. Trends in Genetics. 2010;26: 99–102. doi:10.1016/j.tig.2009.12.012

39. Niknejad A, Comte A, Parmentier G, Roux J, Bastian FB, Robinson-Rechavi M. vHOG, a multispecies vertebrate ontology of homologous organs groups. Bioinformatics. 2012;28: 1017–1020. doi:10.1093/bioinformatics/bts048

40. Franz NM, Goldstein AM. Phenotype ontologies: are homology relations central enough? A reply to Deans et al. Trends in Ecology & Evolution. 2013;28: 131–132. doi:10.1016/j.tree.2012.08.001

41. Owen R. On the Anatomy of Vertebrates. London: Longmans, Green and Co; 1866.

42. Robinson PN, Bauer S. Introduction to Bio-Ontologies. Chapman and Hall/CRC; 2011.

43. OWL 2 Web Ontology Language Profiles (Second Edition) [Internet]. [cited 8 Feb 2019]. Available: https://www.w3.org/TR/owl2-profiles/

44. Mungall CJ, Dietze H, Osumi-Sutherland D. Use of OWL within the Gene Ontology. bioRxiv. 2014; 010090. doi:10.1101/010090

45. Grüninger M, Fox MS. Methodology for the design and evaluation of ontologies. IJCAI’95 Workshop on Basic Ontological Issues in Knowledge Sharing. 1995.

46. Malheiros Y, Freitas F. Unification in EL for competency question generation. Description Logics. 2017. Available: http://ceur-ws.org/Vol-1879/paper42.pdf

47. Mungall C, Gkoutos G, Washington N, Lewis S. Representing phenotypes in OWL. Proceedings of the OWLED 2007 Workshop on OWL. 2007.

48. Mungall CJ, Gkoutos GV, Smith CL, Haendel MA, Lewis SE, Ashburner M. Integrating phenotype ontologies across multiple species. Genome Biology. 2010;11: R2. doi:10.1186/gb-2010-11-1-r2

49. Galis F, van Alphen JJM, Metz JAJ. Why five fingers? Evolutionary constraints on digit numbers. Trends in Ecology & Evolution. 2001;16: 637–646. doi:10.1016/S0169-5347(01)02289-3

50. Fabrezi M. A survey of prepollex and prehallux variation in anuran limbs. Zoological Journal of the Linnean Society. Oxford University Press; 2001;131: 227–248. doi:10.1111/j.1096-3642.2001.tb01316.x

51. Hall BK. Fins into Limbs: Evolution, Development, and Transformation. University of Chicago Press; 2008.

52. Clack JA. Gaining Ground, Second Edition: The Origin and Evolution of Tetrapods. Indiana University Press; 2012.

53. Shou S, Scott V, Reed C, Hitzemann R, Stadler HS. Transcriptome analysis of the murine forelimb and hindlimb autopod. Developmental Dynamics. 2005;234: 74–89. doi:10.1002/dvdy.20514

54. Tamura K, Nomura N, Seki R, Yonei-Tamura S, Yokoyama H. Embryological evidence identifies wing digits in birds as digits 1, 2, and 3. Science. 2011;331: 753–757. doi:10.1126/science.1198229

55. Cranston K, Harmon LJ, O’Leary MA, Lisle C. Best practices for data sharing in phylogenetic research. PLoS Currents. 2014;6. doi:10.1371/currents.tol.bf01eff4a6b60ca4825c69293dc59645

56. Chibucos MC, Mungall CJ, Balakrishnan R, Christie KR, Huntley RP, White O, et al. Standardized description of scientific evidence using the Evidence Ontology (ECO). Database. 2014;2014. doi:10.1093/database/bau075

57. Nelson CE, Morgan BA, Burke AC, Laufer E, DiMambro E, Murtaugh LC, et al. Analysis of Hox gene expression in the chick limb bud. Development. 1996;122: 1449–1466. Available: http://dev.biologists.org/cgi/pmidlookup?view=long&pmid=8625833

58. Ruvinsky I, Gibson-Brown JJ. Genetic and developmental bases of serial homology in vertebrate limb evolution. Development. 2000;127: 5233–5244. Available: http://dev.biologists.org/cgi/pmidlookup?view=long&pmid=11076746

59. Tabin C, Laufer E. Hox genes and serial homology. Nature. 1993;361: 692. doi:10.1038/361692a0

60. Goodrich ES. Studies on the Structure and Development of Vertebrates. London: Macmillan and Co; 1930.

61. Freitas R, Zhang G, Cohn MJ. Biphasic Hoxd gene expression in shark paired fins reveals an ancient origin of the distal limb domain. PLoS One. 2007;2: e754. doi:10.1371/journal.pone.0000754

62. Grau BC, Horrocks I, Kazakov Y, Sattler U. Just the right amount: extracting modules from ontologies. Proceedings of the 16th International Conference on World Wide Web. New York, NY, USA: ACM; 2007. pp. 717–726. doi:10.1145/1242572.1242669

63. Glimm B, Horrocks I, Motik B, Stoilos G, Wang Z. HermiT: An OWL 2 Reasoner. Journal of Automated Reasoning. 2014;53: 245–269. doi:10.1007/s10817-014-9305-1

64. Vargas AO, Fallon JF. Birds have dinosaur wings: The molecular evidence. Journal of Experimental Zoology Part B, Molecular and Developmental Evolution. 2005;304: 86–90. doi:10.1002/jez.b.21023

65. Abouheif E, Akam M, Dickinson WJ, Holland PW, Meyer A, Patel NH, et al. Homology and developmental genes. Trends in Genetics. 1997;13: 432–433. Available: https://www.ncbi.nlm.nih.gov/pubmed/9385839

66. Striedter GF, Northcutt RG. Biological hierarchies and the concept of homology. Brain, Behavior and Evolution. 1991;38: 177–189. doi:10.1159/000114387

67. Roth VL. Within and between organisms: replicators, lineages, and homologues. In: Hall BK, editor. Homology. San Diego: Academic Press; 1994. pp. 301–337. doi:10.1016/B978-0-08-057430-1.50015-9

68. Panganiban G, Irvine SM, Lowe C, Roehl H, Corley LS, Sherbon B, et al. The origin and evolution of animal appendages. Proceedings of the National Academy of Sciences. 1997;94: 5162–5166. doi:10.1073/pnas.94.10.5162

69. Shubin N, Tabin C, Carroll S. Fossils, genes and the evolution of animal limbs. Nature. 1997;388: 639–648. doi:10.1038/41710

70. Shubin N, Tabin C, Carroll S. Deep homology and the origins of evolutionary novelty. Nature. 2009;457: 818–823. doi:10.1038/nature07891

71. Kazakov Y, Krötzsch M, Simančík F. The Incredible ELK. Journal of Automated Reasoning. 2014;53: 1–61. doi:10.1007/s10817-013-9296-3

72. Tarasov S. Integration of anatomy ontologies and evo-devo using structured Markov models suggests a new framework for modeling discrete phenotypic traits. bioRxiv preprint. 2018; doi:10.1101/188672.

73. Chibucos MC, Siegele DA, Hu JC, Giglio M. The Evidence and Conclusion Ontology (ECO): supporting GO annotations. In: Dessimoz C, Škunca N, editors. The Gene Ontology Handbook. New York, NY: Springer New York; 2017. pp. 245–259. doi:10.1007/978-1-4939-3743-1_18

74. Giglio M, Tauber R, Nadendla S, Munro J, Olley D, Ball S, et al. ECO, the Evidence & Conclusion Ontology: community standard for evidence information. Nucleic Acids Research. 2019;47: D1186–D1194. doi:10.1093/nar/gky1036

75. Hiller M, Schaar BT, Indjeian VB, Kingsley DM, Hagey LR, Bejerano G. A “forward genomics” approach links genotype to phenotype using independent phenotypic losses among related species. Cell Reports. 2012;2: 817–823. doi:10.1016/j.celrep.2012.08.032

76. Lippert C, Sabatini R, Maher MC, Kang EY, Lee S, Arikan O, et al. Identification of individuals by trait prediction using whole-genome sequencing data. Proceedings of the National Academy of Sciences. 2017;114: 10166–10171. doi:10.1073/pnas.1711125114

