## Supplementary Materials 3 for "A logical model of homology for comparative biology"

Supplementary Materials 3. OWL explanation for 'length of forelimb' as phenotype of a 'pectoral fin' homolog. The explanation does not require evolutionary "grouping classes" found in Uberon, such as 'paired limb/fin'.

  length#q3 'inheres in' forelimb#e3
  'forelimb' SubClassOf 'limb'
  'limb' SubClassOf only_in_taxon some 'Tetrapoda'
  forelimb#e3 Type 'forelimb'
  only_in_taxon SubPropertyOf in_taxon
  'forelimb' and (in_taxon some Tetrapoda) SubClassOf in_historical_homology_relationship_with some ('pectoral fin' and (in_taxon some 'Vertebrata'))
  Question1 EquivalentTo inheres_in some (in_historical_homology_relationship_with some 'pectoral fin')
